# Age-Related Differences in Human Cortical Microstructure Depend on the Distance to the Nearest Vein

**DOI:** 10.1101/2023.12.20.572529

**Authors:** Christoph Knoll, Juliane Doehler, Alicia Northall, Stefanie Schreiber, Johanna Rotta, Hendrik Mattern, Esther Kuehn

## Abstract

Age-related differences in cortical microstructure are used to understand the neuronal mechanisms that underlie human brain ageing. The cerebral vasculature contributes to cortical ageing, but its precise interaction with cortical microstructure is poorly understood. In a cross-sectional study, we combine venous imaging with vessel distance mapping (VDM) to investigate the interaction between venous distances and age-related differences in the microstructural architecture of the primary somatosensory cortex (S1), the primary motor cortex (M1), and additional areas in frontal cortex as non-sensorimotor control regions. We scanned 18 younger adults and 17 older adults using 7T-MRI to measure age-related changes in quantitative T1 (qT1) values, positive QSM (pQSM) values, and negative QSM (nQSM) values at 0.5 mm isotropic resolution. We modelled different cortical depths using an equi-volume approach and assessed the distance of each voxel to its nearest vein using VDM. Our data reveal a dependence of cortical qT1 and pQSM values on venous distance. In addition, there is an interaction between venous distance and age on qT1 values, driven by lower qT1 values in older compared to younger adults in voxels that are closer to a vein. Together, our data show that the local venous architecture explains a significant amount of variance in standard measures of cortical microstructure and should be considered in neurobiological models of human brain organisation and cortical ageing.

## Introduction

Age-related differences in cortical microstructure are used to understand the neuronal mechanisms that underlie human brain ageing and neurodegeneration. Vascular supply patterns in the brain vary between individuals^1,2^, interact with cortical microstructure and functional decline^3,4^, and contribute to brain pathology.^5^ However, the interaction between the local vascular architecture and age-related changes in cortical microstructure is poorly understood. This knowledge gap prevents us from understanding the mechanisms that underlie human cortical ageing and potential protective factors.

The human cortex is permeated by a dense net of blood vessels that is highly variable in location and diameter.^2^ Vascular changes are a risk factor for all-cause dementia^6^, and antihypertensive treatment can lower dementia incidence.^7^ A human MRI study reported that participants with cerebral small vessel disease, who show a double-supply pattern of hippocampal vascularization, perform better in cognitive tests compared to those with a single-supply pattern.^3^ Vascular health is regarded as a key driver for resilience in ageing, for example by maintaining blood flow and neuronal metabolism.^8^

However, it is still unclear how individual vascular supply patterns interact with age-related cortical degeneration. Cortical locations that are more distant to larger blood vessels might be more prone to age-related structural alterations, such as myelin loss^9–11^ and iron accumulation^10,12–14^, in case ageing blood vessels reduce supply to locations farther away from major branches (Hypothesis 1, H1). In this view, cortical locations that are closer to a vein may be more protected from age-related decline. Alternatively, myelin loss and iron accumulation might be more pronounced at cortical locations that are closer to blood vessels, because age-related substance accumulation and calcification occur near or within blood vessels^15^ (Hypothesis 2, H2). In this view, cortical locations that are farther away from a vein may be more protected from age-related decline. Finally, cortical degeneration may occur independently of the distance to blood vessels (Hypothesis 3, H3).

To test these hypotheses, we compared two extreme age groups (healthy younger adults < 30 years, healthy older adults > 65 years) using ultra-high resolution magnetic resonance imaging at 7 Tesla (7T-MRI). Quantitative T1 values (qT1) were used as validated proxy for cortical myelin content.^16–18^ Quantitative susceptibility mapping (QSM) was used to quantify cortical iron^12,13,19–21^ (positive QSM, pQSM) and calcium/proteins^20,22–26^ (negative QSM, nQSM). We focus on primary somatosensory cortex (S1) and primary motor cortex (M1) as both areas play an important role in ageing and neurodegeneration, and venous density in S1 and M1 has been associated with myelin loss and declined sensorimotor function in older adults.^5^ Investigations of S1 and M1 microstructure remain rare, but may explain functional loss in hands and feet in older age, which may impact daily life routines.^27^ In addition, we investigate several areas in the frontal cortex (superior frontal gyrus (SFG), caudal middle frontal cortex (CMF), rostral middle frontal cortex (RMF)) as non-sensorimotor control regions (results shown in **Supplementary Material**). QSM images were used for vein identification. Vessel Distance Mapping (VDM) was used to link age-related changes in cortical microstructure to the distance to the nearest vein.^28,29^

We combine MRI proxies of cortical myelin, iron and calcium/proteins with VDM to test whether age-related changes in S1 and M1 microstructure show a relation to the distance to the nearest vein. This work contributes to our understanding of how the individual vascular architecture relates to microstructural variance in the ageing cortex, and provides critical insights into the mechanisms of cortical ageing.

## Materials & Methods

### Participants

35 healthy volunteers (n=18 younger adults: M=25 years, SD=3 years, range=21-29 years, 9 females; n=17 older adults: M=70 years, SD=4 years, range=65-77 years, 10 females) were recruited from the database of the German Center for Neurodegenerative Diseases (DZNE), Magdeburg, Germany and participated in our cross-sectional study. All participants were right-handed (laterality index ranging from +40 to +100).^30^

7T-MRI contraindications, chronic illness, psychiatric and neurological disorders, central acting medications and impaired hand function were exclusion criteria. 1/17 older adults underwent carpal tunnel surgery on both hands. However, no complaints were reported and no outliers were detected in sensorimotor tasks (see **Supplementary Table 1**).

Regarding vascular health, 6/17 older adults reported a history of hypertension that was well-controlled with specific medication in all cases. Two experienced neurologists evaluated vascular health of older adults according to the STandards for ReportIng Vascular changes on nEuroimaging (STRIVE-2)^31,32^ and the guideline by Chen et al.^33^ We detected microbleeds, cortical superficial siderosis, lacunae and/or enlarged perivascular spaces in 6/17 older adults (see **Table 1**). All anomalies were detected outside S1 and M1. In total, 9/17 older adults showed vascular health risk factors (hypertension and/or signs of cerebral small vessel diseases). Besides, participants showed no medical illnesses. As indicated by the ‘Montreal Cognitive Assessment’ (MoCA)^34^, there were no signs of cognitive impairments (mean score 28.33 ± 1.45), except for one older adult with a MoCA score of 21 (MoCA ≥ 26 is considered normal^34^). Because this participant performed equally well in sensorimotor tasks compared with all other older adults (see **Supplementary Table 1**), we kept the data in the main analysis and performed additional control analyses excluding this participant. There were no professional musicians among the participants.^35–37^

**Table 1.** MoCA scores and vascular health rating for older adults. The ‘Montreal Cognitive Assessment’ (MoCA) score is an indicator of cognitive impairment. A MoCA score ≥ 26 is considered normal. Vascular health is given as self-reported history of hypertension (y=yes, n=no), visually detected thrombosis following the guideline by Chen et al.^33^ and the STRIVE-2 criteria^32^ (cSS=cortical superficial siderosis; EPVS=enlarged perivascular spaces). In parentheses, the number of lesions are detailed. The composite score reflects the sum of positive STRIVE-2 criteria (1=no positive criterion, 2=one positive criteria, 3=two positive criteria, 4=three positive criteria).

| Participant | MoCA | Hypertension | Thrombosis | STRIVE-2 Criteria |  |  |  |  | Composite Score |
| --- | --- | --- | --- | --- | --- | --- | --- | --- | --- |
|  |  |  |  | Cerebral Microbleeds | cSS | Microinfarcts | Lacunae | EPVS |  |
| 1 | 26 | n | n | n | n | n | n | n | 1 |
| 2 | 21 | n | n | n | n | n | n | n | 1 |
| 3 | 29 | y | n | n | n | n | subcortical frontal L (1) | n | 2 |
| 4 | 28 | n | n | n | n | n | n | n | 1 |
| 5 | 28 | n | n | temporal R (1) | frontal R | n | frontal L, parietal L+R (3) | n | 4 |
| 6 | 29 | n | n | n | n | n | n | n | 1 |
| 7 | 30 | y | n | n | n | n | deep white matter L+R (3) | n | 2 |
| 8 | 26 | n | n | n | n | n | n | multiple | 2 |
| 9 | 27 | y | n | n | n | n | n | n | 1 |
| 10 | 28 | y | n | n | n | n | n | n | 1 |
| 11 | 26 | n | n | temporal R (1) | n | n | n | n | 2 |
| 12 | 26 | n | n | n | n | n | n | n | 1 |
| 13 | - | y | n | n | n | n | n | n | 1 |
| 14 | 28 | n | n | n | n | n | n | n | 1 |
| 15 | 29 | y | n | occipital R (1) | parietal R,<br>frontal L+R | n | n | multiple | 4 |
| 16 | 26 | n | n | n | n | n | n | n | 1 |
| 17 | 30 | n | n | n | n | n | n | n | 1 |

All participants were paid for their attendance and provided written informed consent. The study was approved by the local Ethics committee of the Otto-von-Guericke University Magdeburg, Germany. Structural S1 and M1 data was published previously^14,38^. However, the venous architecture was not investigated before.

### MRI Data Acquisition

All participants took part in one structural 7T-MRI session. Data was acquired at a Siemens MAGNETOM scanner located in Magdeburg, Germany, using a 32-channel head coil. First, MP2RAGE^39^ whole-brain images were acquired at 0.7 mm isotropic resolution (240 sagittal slices, field-of-view=224 mm, repetition time=4800 ms, echo time=2.01 ms, inversion time TI1/TI2=900/2750 ms, flip angle=5◦/3◦, bandwidth=250 Hz/Px, GRAPPA 2). Second, MP2RAGE part-brain images (optimised for M1 and S1) were acquired at 0.5 mm isotropic resolution (208 transversal slices, field-of-view read=224 mm, repetition time=4800 ms, echo time=2.62 ms, inversion time TI1/TI2=900/2750 ms, flip angle=5◦/3◦, bandwidth=250 Hz/Px, GRAPPA 2, slab oversampling=7.7 %). Finally, part-brain susceptibility-weighted images (SWI) were acquired at 0.5 mm isotropic resolution using a 3D GRE sequence^40^ (208 transversal slices, field-of-view read=192 mm, repetition time=22 ms, echo time=9.00 ms, flip angle=10◦, bandwidth=160 Hz/Px, GRAPPA 2, slice oversampling=7.7 %). The total scanning time was around 60 minutes.

### Image Preprocessing

We used the same preprocessing pipeline as described previously.^14,38^ In short, data quality was ensured by two independent raters. QSM images were reconstructed using the Bayesian multiscale dipole inversion (MSDI) algorithm (qsmbox v2.0, https://gitlab.com/acostaj/QSMbox).^41^ MP2RAGE and non-normalized QSM data^13^ were processed using JIST^42^ and CBS Tools^43^ as plug-ins for MIPAV (https://mipav.cit.nih.gov/).

Slab qT1 images were registered to upsampled whole-brain qT1 images (ANTs v1.9x embedded in CBS Tools)^44^ in one single step before slab and whole-brain images were merged. Extra-cranial tissue and dura mater were removed^43^. QSM images were registered to the merged whole-brain qT1 images using ITK-SNAP (v3.6.0, www.itksnap.org).

Cortical segmentation was calculated on the UNI image, using the TOADS algorithm.^45^ To estimate the boundaries between white matter (WM) and grey matter (GM), and between GM and cerebrospinal fluid, the CRUISE algorithm^46^ was used. Resulting level set surfaces^47^ were optimised to precisely match M1 and S1 (based on empirical optimization).^14,38^

The cortex was divided into 21 cortical depths using the validated equi-volume model.^48^ After removing the three most superficial and the two deepest cortical depths to reduce partial volume effects^49,50^, the remaining 16 depths were averaged into four equally-spaced compartments (SF=superficial, OM=outer-middle, IM=inner-middle, DP=deep).^49^ The resulting cortical depths do not correspond to anatomical layers. Finally, the non-merged high-resolution qT1 and QSM slab data were used to extract qT1 values, positive QSM (pQSM) values, and negative QSM (nQSM) values.

### Vessel Segmentation and Vessel Distance Mapping (VDM)

The Filtered Vessels module (v3.0.4; CBS Tools) was used to extract vessels from the QSMs.^43^ The vasculature was segmented using a Markov Random Field diffusion technique. Previous validation studies revealed that this technique captures venous architectures with high precision.^51^ Resulting vessel probability maps represent the probability of each voxel belonging to a vessel.

For computing VDMs^29^, individual vessel probability maps were binarized. For thresholding, we used the Openly available sMall vEsseL sEgmenTa-Tion pipelinE (OMELETTE, https://gitlab.com/hmattern/omelette).^52^ Thresholds were set using a 3-class Otsu histogram analysis (returning a lower and upper threshold). Voxels above the upper threshold were considered as vessels, voxels below the lower threshold were considered as background. To classify vessels between the lower and upper threshold, hysteresis thresholding was used^53^, which considers a voxel belonging to a vessel only if it is connected to a voxel with a value above the upper threshold. A Euclidean distance transformation^54^ was applied to the thresholded data to compute the distance to the closest segmented vein per voxel. Resulting VDMs were multiplied with binarized cortical depth masks and brain area masks (S1, M1, SFG, CMF, RMF), generating one VDM per cortical depth, brain area and participant (see **Figure 1**). qT1, pQSM and nQSM images were multiplied with inverted binarized vessel probability masks (extracted values not overlapping with veins).

**Figure 1:**
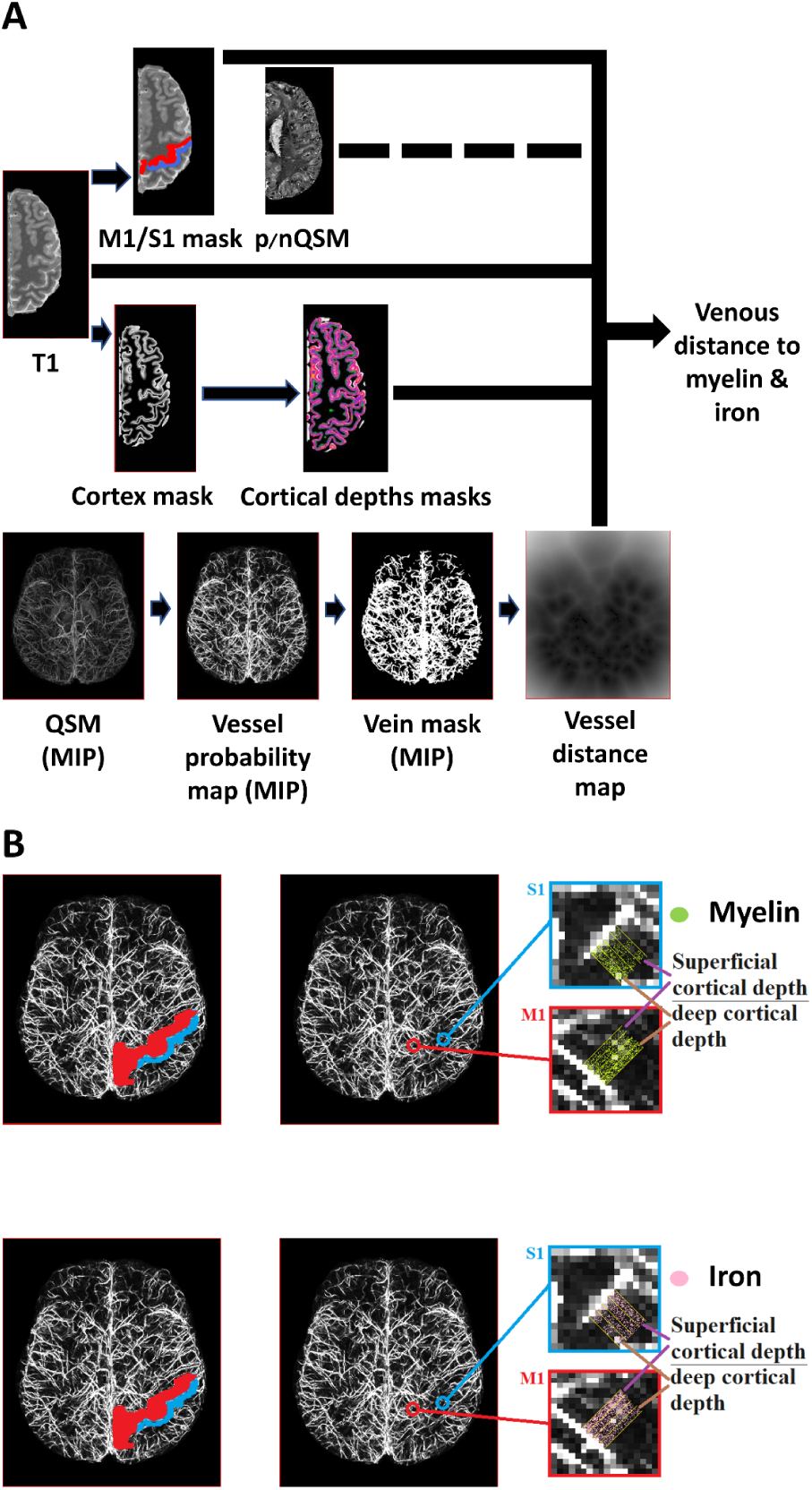
Overview Methodology and Analysis. **A:** qT1 and QSM images were sampled at different cortical depths (cortical depths masks) after cortex segmentation (cortex masks). Masks were applied to extract data from the primary motor cortex (M1, *red mask*) and primary somatosensory cortex (S1, *blue mask*). As a non-sensorimotor control region, data was extracted from the prefrontal cortex (SFG, CMF, RMF, not shown). Quantitative Susceptibility Maps (QSM; displayed as Maximum Intensity Projections, MIP) were used to extract vessel probability maps (including intensity values between 0 and 1) to identify venous vasculature. A hysteresis filter was used to generate a binarized vein mask before a Euclidean distance transformation was applied to compute the Vessel Distance Map (VDM). Individual VDMs were multiplied with binarized cortical depth masks and then applied to the qT1 image before resulting parameter maps were multiplied with binarized M1/S1/SFG/CMF/RMF masks (*analysis pathway shown as black lines*). Please note that the same analysis pathway was applied to the pQSM and nQSM data (*black dotted line*). **B:** Shown are masks covering left M1 (*red*) and left S1 (*blue*) together with the vessel probability map of one example participant. Magnified images show extracted cortical depth compartments in relation to the distance to the nearest vein.

### Definition of Brain Areas

M1 and S1 masks were manually generated using anatomical landmarks^55–58^, following a standardised procedure^14,38,59,60^. We masked all slices in M1 in which the hand knob was visible^56^, and extended the masks until the precentral gyrus was completely covered.^14,59^ S1 masks were drawn from the crown of the postcentral gyrus to the fundus of the central sulcus^38^, covering area 3b and parts of areas 1 and 3a.^55^ M1 masks also include medial M1, whereas S1 masks mainly cover lateral S1. Resulting masks were plotted in reference to co-registered Freesurfer labels to ensure overlap with automated approaches.

The masks for SFG, CMF, RMF were generated using Freesurfer (v7.4.1). Whole-brain T1-weighted images (0.7 mm isotropic) were used as input for the recon-all command to segment the cortex based on the Desikan-Killiany atlas^61^ in individual subject space. Resulting labels were upsampled and registered to the slab images (0.5 mm isotropic). Binarized labels were used as SFG, CMF and RMF masks.

### Distance Binning

Each cortical VDM was discretized into 5 bins that represent the distance to the nearest vein (bin 0-2: 0.01–2 mm distance to the nearest vein, bin 2-4: 2.01–4 mm, bin 4-6: 4.01–6 mm, bin 6-8: 6.01–8 mm, bin 8-10: 8.01–10 mm). This was calculated for each voxel in each brain area. Binning was used to obtain different distance conditions for statistical analysis. We chose 5 bins to include a minimum of 4 voxels per distance condition. Resulting binned data (5 VDMs per cortical depth and brain region) was used to extract qT1, pQSM and nQSM values.

### Vascular Health Evaluation

Two experienced neurologists evaluated vascular health of older adults based on visual inspection of MR scans. Following the guideline by Chen et al.^33^, thromboses were investigated using part-brain QSM and SWI data. According to the STRIVE-2 recommendations, microbleeds and cortical superficial siderosis were investigated based on part-brain SWI data, whereas lacunae and microinfarcts were examined using whole-brain T1-weighted images.

### Behavioural Tasks

Outside of the MRI scanner, tactile detection thresholds were assessed using fine hair stimuli, tactile 2-point discrimination performance was assessed using a standard procedure^62–64^, and sensorimotor integration performance was assessed with a custom made pressure sensor^65^ (see **Supplementary Methods** for details).

### Statistics

To test for a main effect of venous distance and an interaction between venous distance and age on cortical microstructure, we calculated mixed-effects ANOVAs on qT1, pQSM and nQSM values with age group (younger adults, older adults) as between-subjects factor, and brain area (M1, S1), cortical depth (SF, OM, IM, DP) and distance to the nearest vein (bin 0-2, bin 2-4, bin 4-6, bin 6-8, bin 8-10; the same excluding bin 0-2) as within-subjects factors. The same mixed-effects ANOVAs were calculated with the SFG, CMF and RMF included as additional brain areas; results of these additional analyses are shown in the **Supplementary Material**. Statistical analyses were performed using SPSS (v21.0.) and R (v4.2.2). Sample distributions were tested for normality using Shapiro-Wilk’s test in combination with visual inspections (see **Supplementary Figures 1, 2**). Homogeneity of variances was tested with Levene’s test. In case of sphericity violations, Greenhouse-Geisser-corrected results were used. Post-hoc tests were performed as two-tailed paired-samples *t*-tests to follow up within-subjects effects and as two-tailed independent samples Welch *t*-tests to follow up between-subjects effects. Given the bin representing the closest distance to the nearest vein (bin 0-2) contained no qT1 values in 12/18 younger adults and 4/17 older adults, no pQSM values in 11/18 younger and 8/17 older adults, and no nQSM values in 13/18 younger and 8/17 older adults, we report statistical analyses both for 5 distance conditions with imputed missing values using the method of multiple monotone imputation based on linear regression, and for 4 distance conditions (excluding bin 0-2). The significance level was set to p≤0.05. In addition to uncorrected p-values, we report Holm-Bonferroni corrected p-values. Generalised Eta-squared (η²_G_) was calculated as an effect size estimator for ANOVAs.^66^ Cohen’s benchmarks of 0.06 and 0.14 for medium and large effects, respectively, were applied.^67^ Hedges-corrected Cohen’s d was calculated as an effects size measure for post-hoc t-tests^68^ to account for small sample sizes (rstatix package v0.7.2). Confidence intervals for Cohen’s d were bootstrapped with the percentile method.^69^ Cohen’s benchmarks of 0.2, 0.5 and 0.8 for small, moderate and large effects, respectively, were applied .^67^

To control for a possible effect of vascular health risk on qT1, pQSM and nQSM values in older adults, we calculated random intercept models using the lmer function (lme 4 package v1.1.33). We compared two models (model 1: structural measure ∼ distance to nearest vein + vascular risk + 1/participant; model 2: structural measure ∼ distance to nearest vein + vascular risk + distance to nearest vein:vascular risk + 1/participant) with a null model (structural measure ∼ distance to nearest vein + 1/participant).

Permutation Welch two sample t-tests were calculated using the perm.t.test function (MKinfer package v1.1) with a number of 100,000 Monte-Carlo permutations. Finally, we correlated pQSM and qT1 values for all area-by-distance conditions separately for younger and older adults using Pearson correlation coefficients (misty package v0.6.2).

## Results

### Age-related Differences in Cortical Myelination Interact with the Distance to the Nearest Vein

To test for the main effect of venous distance on qT1 values and a possible interaction with age, we computed an ANOVA with the factors age (younger adults, older adults), brain area (S1, M1), cortical depth (SF, OM, IM, DP) and venous distance (bin 0-2, bin 2-4, bin 4-6, bin 6-8, bin 8-10) on qT1 values. As expected, there is a significant main effect of brain area driven by lower qT1 values (higher myelination) in M1 compared to S1, and a significant main effect of cortical depth driven by lower qT1 values (higher myelination) in deeper compared to superficial depths in M1 and S1 (see **Table 2**, **Figure 2**).

**Figure 2.**
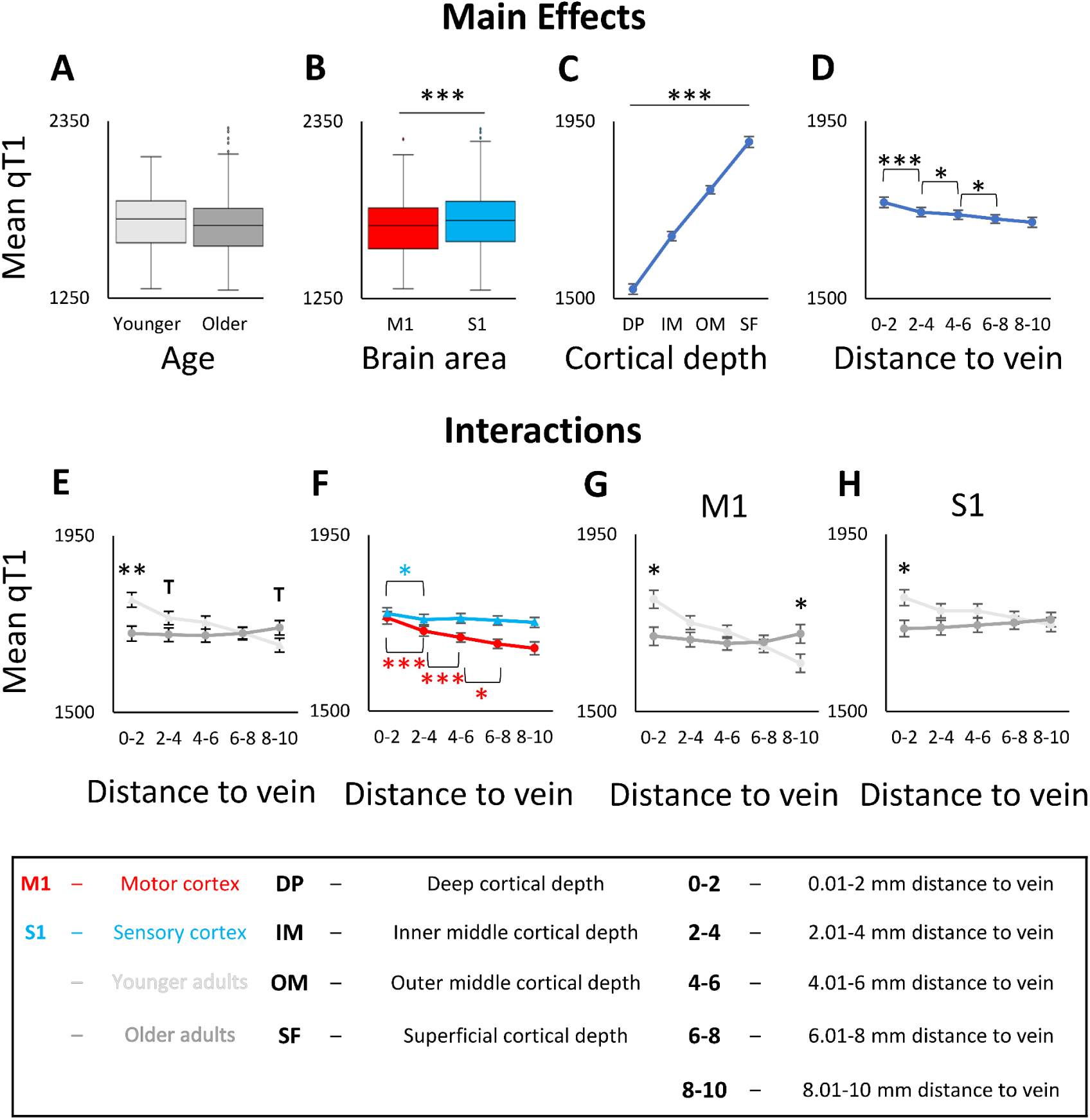
Interaction between age, brain area, cortical depth and venous distance for cortical qT1 values. qT1 values are given in milliseconds (lower values indicate higher myelin content). **A:** No significant main effect of age on qT1 values. Shown are medians, interquartile ranges, and lower and upper quartiles for younger (n=18, *light grey*) and older (n=17, *dark grey*) adults. Dots above a box mark outliers. **B:** Significant main effect of brain area (i.e., M1 (*red*), S1 (*blue*)) on qT1 values. **C:** Significant main effect of cortical depth on qT1 values; dots represent mean qT1 values for the different cortical depths averaged across age groups, distances, and brain areas. Error bars show standard errors of the mean (SEM). **D:** Significant main effect of venous distance on qT1 values (0-2 = 0.01-2 mm, 2-4 = 2.01-4 mm, 4-6 = 4.01-6 mm, 6-8 = 6.01-8 mm, 8-10 = 8.01-10 mm). **E:** Significant interaction effect between venous distance and age on qT1 values. **F:** Significant interaction effect between venous distance and brain area on qT1 values. **G/H:** Significant interaction effect between venous distance, age and brain area (**G**: data for M1, **H**: data for S1). Significant results of post-hoc t-tests to follow-up mixed-effects ANOVA results (see **Table 2** for statistical results) are marked by asterisks: * p≤0.05, ** p≤0.005, *** p≤0.0005 (uncorrected). Trends above p=0.05 are marked by a T.

**Table 2.** ANOVA Results for qT1 values. Results for the mixed-effects ANOVA on qT1 values with factors age (n=18 younger adults, n=17 older adults), cortical depth (SF, OM, IM, DP), brain area (S1, M1) and venous distance (5 distances). Given are mean qT1 values in milliseconds (Mean in ms) and standard errors of the mean (SEM), degrees of freedom of the numerator (DFn), degrees of freedom of the denominator (DFd), p-values (p), and effect sizes (n_G_²). P-values ≤ 0.05 are considered as significant.

| Effect | Conditions | Mean<br>(ms) | SEM<br>(ms) | Statistics |  |  |  |  |
| --- | --- | --- | --- | --- | --- | --- | --- | --- |
| | | | | DFn | DFd | F | p | $\eta_G^2$ |
| Age | Younger | 1725.97 | 15.94 | 1 | 33 | 1.092 | 0.304 | 0.018 |
|  | Older | 1702.08 | 16.40 |  |  |  |  |  |
| Brain Area | M1 | 1692.42 | 12.71 | 1 | 33 | 24.856 | 1.924*10<sup>-5</sup> | 0.056 |
|  | S1 | 1735.63 | 11.73 |  |  |  |  |  |
| Cortical depth | Deep | 1523.86 | 13.13 | 3 | 31 | 909.332 | 2.285*10<sup>-37</sup> | 0.708 |
|  | Inner Middle | 1658.11 | 11.65 |  |  |  |  |  |
|  | Outer Middle | 1776.34 | 10.22 |  |  |  |  |  |
|  | Superficial | 1897.80 | 13.98 |  |  |  |  |  |
| Distance | 0.01 - 2 mm | 1743.83 | 13.46 | 4 | 30 | 16.289 | 9.651*10<sup>-6</sup> | 0.036 |
|  | 2.01 - 4 mm | 1719.03 | 12.16 |  |  |  |  |  |
|  | 4.01 - 6 mm | 1712.38 | 11.53 |  |  |  |  |  |
|  | 6.01 - 8 mm | 1701.47 | 10.89 |  |  |  |  |  |
|  | 8.01 - 10 mm | 1693.41 | 12.87 |  |  |  |  |  |
| Age x Distance | Younger / 0.01 - 2 mm | 1786.91 | 18.76 | 4 | 30 | 25.657 | 7.668*10<sup>-8</sup> | 0.056 |
|  | Younger / 2.01 - 4 mm | 1740.50 | 16.95 |  |  |  |  |  |
|  | Younger / 4.01 - 6 mm | 1728.96 | 16.06 |  |  |  |  |  |
|  | Younger / 6.01 - 8 mm | 1702.08 | 15.17 |  |  |  |  |  |
|  | Younger / 8.01 - 10 mm | 1671.42 | 17.94 |  |  |  |  |  |
|  | Older / 0.01 - 2 mm | 1700.76 | 19.31 |  |  |  |  |  |
|  | Older / 2.01 - 4 mm | 1697.57 | 17.44 |  |  |  |  |  |
|  | Older / 4.01 - 6 mm | 1695.80 | 16.53 |  |  |  |  |  |
|  | Older / 6.01 - 8 mm | 1700.87 | 15.61 |  |  |  |  |  |
|  | Older / 8.01 - 10 mm | 1715.39 | 18.46 |  |  |  |  |  |
| Age x Brain Area | Younger / M1 | 1700.53 | 17.71 | 1 | 33 | 0.783 | 0.382 | 0.002 |
|  | Older / M1 | 1684.31 | 18.23 |  |  |  |  |  |
|  | Younger / S1 | 1751.41 | 16.36 |  |  |  |  |  |
|  | Older / S1 | 1719.85 | 16.83 |  |  |  |  |  |
| Brain Area x Distance | M1 / 0.01 - 2 mm | 1738.28 | 16.18 | 4 | 30 | 8.628 | 2.325*10<sup>-4</sup> | 0.013 |
|  | M1 / 2.01 - 4 mm | 1704.16 | 12.67 |  |  |  |  |  |
|  | M1 / 4.01 - 6 mm | 1687.94 | 11.69 |  |  |  |  |  |
|  | M1 / 6.01 - 8 mm | 1671.56 | 11.81 |  |  |  |  |  |
|  | M1 / 8.01 - 10 mm | 1660.16 | 16.50 |  |  |  |  |  |
|  | S1 / 0.01 - 2 mm | 1749.39 | 14.49 |  |  |  |  |  |
|  | S1 / 2.01 - 4 mm | 1733.91 | 12.96 |  |  |  |  |  |
|  | S1 / 4.01 - 6 mm | 1736.82 | 12.58 |  |  |  |  |  |
|  | S1 / 6.01 - 8 mm | 1731.39 | 11.47 |  |  |  |  |  |
|  | S1 / 8.01 - 10 mm | 1726.65 | 12.68 |  |  |  |  |  |
| Age x Brain Area x Distance | Younger / M1 / 0.01 - 2 mm | 1785.29 | 22.55 | 4 | 30 | 3.714 | 0.024 | 0.006 |
|  | Younger / M1 / 2.01 - 4 mm | 1725.63 | 17.66 |  |  |  |  |  |
|  | Younger / M1 / 4.01 - 6 mm | 1702.89 | 16.30 |  |  |  |  |  |
|  | Younger / M1 / 6.01 - 8 mm | 1666.23 | 16.46 |  |  |  |  |  |
|  | Younger / M1 / 8.01 - 10 mm | 1622.62 | 23.00 |  |  |  |  |  |
|  | Younger / S1 / 0.01 - 2 mm | 1788.53 | 20.20 |  |  |  |  |  |
|  | Younger / S1 / 2.01 - 4 mm | 1755.36 | 18.06 |  |  |  |  |  |
|  | Younger / S1 / 4.01 - 6 mm | 1755.03 | 17.53 |  |  |  |  |  |
|  | Younger / S1 / 6.01 - 8 mm | 1737.92 | 15.99 |  |  |  |  |  |
|  | Younger / S1 / 8.01 - 10 mm | 1720.23 | 17.68 |  |  |  |  |  |
|  | Older / M1 / 0.01 - 2 mm | 1691.27 | 23.21 |  |  |  |  |  |
|  | Older / M1 / 2.01 - 4 mm | 1682.68 | 18.17 |  |  |  |  |  |
|  | Older / M1 / 4.01 - 6 mm | 1672.99 | 16.77 |  |  |  |  |  |
|  | Older / M1 / 6.01 - 8 mm | 1676.88 | 16.94 |  |  |  |  |  |
|  | Older / M1 / 8.01 - 10 mm | 1697.71 | 23.66 |  |  |  |  |  |
|  | Older / S1 / 0.01 - 2 mm | 1710.25 | 20.79 |  |  |  |  |  |
|  | Older / S1 / 2.01 - 4 mm | 1712.45 | 18.58 |  |  |  |  |  |
|  | Older / S1 / 4.01 - 6 mm | 1718.60 | 18.04 |  |  |  |  |  |
|  | Older / S1 / 6.01 - 8 mm | 1724.86 | 16.45 |  |  |  |  |  |
|  | Older / S1 / 8.01 - 10 mm | 1733.07 | 18.19 |  |  |  |  |  |

There is also a significant main effect of venous distance driven by significantly higher qT1 values (reduced myelination) closer compared to farer away from a vein (significant between bin 0-2 and bin 2-4 (t(34)=4.428, p=9.353*10^-5^, p_Holm-Bonferroni_=3.74*10^-4^, d=0.74), between bin 2-4 and bin 4-6 (t(34)=2.710, p=0.010, p_Holm-Bonferroni_=0.03, d=0.43), and between bin 4-6 and bin 6-8 (t(34)=2.371, p=0.024, p_Holm-Bonferroni_=0.048, d=0.38; see **Table 2**, **Figure 2D**). The venous distance explains 3.6% of the variance in the cortical qT1 signal.

The significant interaction between brain area and venous distance (see **Table 2**) reveals that this effect is mainly driven by M1 rather than by S1. Post hoc *t*-tests show that in M1, qT1 values are significantly higher in bin 0-2 compared to bin 2-4 (t(34)=4.190, p=1.871*10^-5^, p_Holm-Bonferroni_=7.484*10^-5^, d=0.69), in bin 2-4 compared to bin 4-6 (t(34)=4.443, p=8.952*10^-5^, p_Holm-Bonferroni_=2.686*10^-4^, d=0.73), and in bin 4-6 compared to bin 6-8 (t(34)=2.640, p=0.012, p_Holm-Bonferroni_=0.024, d=0.45). In S1, qT1 values are only higher in bin 0-2 compared to bin 2-4, however, this result does not survive Holm-Bonferroni correction (t(34)=2.402, p=0.022, p_Holm-Bonferroni_=0.088, d=0.40). All other comparisons are not significant (bin 2-4 versus bin 4-6: t(34)=-0.823, p=0.416, d=-0.14; bin 4-6 versus 6-8: t(34)=1.063, p=0.295, d=0.18; bin 6-8 versus bin 8-10: t(34)=0.790, p=0.435, d=0.13).

We then explored potential age effects and detected no significant main effect of age and no significant interaction between age and brain area (see **Table 2**, **Figure 2**). However, there is a significant interaction between age and venous distance on qT1 values (see **Table 2**). Post hoc *t*-tests reveal significantly lower qT1 values (higher myelination) in older adults compared to younger adults in bin 0-2 (t(23.345)=-3.145, p=0.004, p_Holm-Bonferroni_=0.02, d=-1.05) and a trend in the same direction in bin 2-4 (t(22.878)=-1.756, p=0.092, d=-0.58), but a reversed trend in bin 8-10 (t(26.499)=1.745, p=0.093, d=0.56; see **Figure 2E**).

Finally, a significant three-way interaction between age, brain area and venous distance (see **Table 2**) reveals that the interaction between age and venous distance is mainly driven by M1 rather than by S1 (see also **Figure 3**). In M1, post hoc *t*-tests reveal significantly lower qT1 values (higher myelination) in older compared to younger adults in bin 0-2 (t(21.015)=-2.845, p=0.010, p_Holm-Bonferroni_=0.05, d=-0.95), and the reverse effect in bin 8-10 (t(33)=2.276, p=0.029, p_Holm-Bonferroni_=0.116, d=0.75). The latter does not survive Holm-Bonferroni correction, but presents with a moderate effect size (see **Figure 3**). In S1, post hoc *t*-tests reveal significantly lower qT1 values (higher myelination) in older compared to younger adults in bin 0-2 (t(33)=-2.701, p=0.011, p_Holm-Bonferroni_=0.05, d=-0.89, see **Figure 2G,H**), but not at other distances. These results are confirmed by permutation t-tests (see **Supplementary Table 2**).

**Figure 3.**
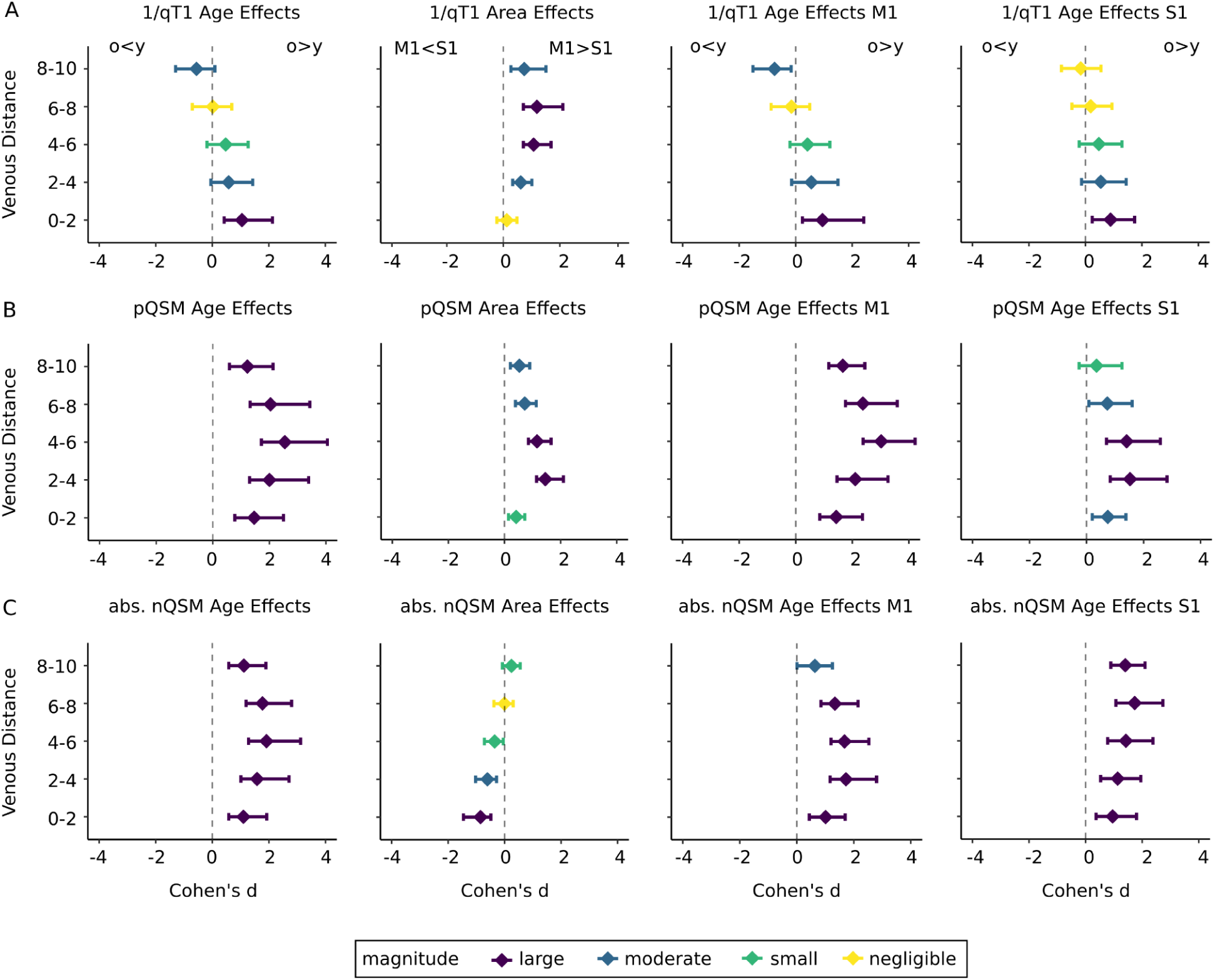
Effect Sizes for the effects of age and brain area on qT1, pQSM and nQSM values plotted in relation to venous distance. Effect sizes (coloured dots) are given as Cohen’s d (standardised difference between group means), venous distances are given in millimetres. qT1 and nQSM values were reversed before effect size calculations, so that larger effect sizes always indicate higher substance in older compared to younger adults (o>y) and in M1 compared to S1 (M1>S1). Different colours indicate the magnitude of the effect sizes (large effects: *purple*, moderate effects: *blue*, small effects: *green*, negligible effects: *yellow*). Horizontal lines indicate 95% confidence intervals. **A**: Effect sizes shown for 1/qT1 values. From left to right: age effects averaged across areas (S1 and M1), area effects averaged across age groups (younger and older adults), age effects for M1, age effects for S1. **B**: Same as in A but effect sizes shown for pQSM values. **C**: Same as in A but effect sizes shown for absolute (abs.) nQSM values.

The interaction effects remain significant when excluding the closest bin (0-2, see **Supplementary Table 3**) and when removing one older participant with a MoCA score below the cut-off for healthy ageing (see **Supplementary Table 4**).

Controlling for effects of vascular health, comparisons of random intercept models reveal no significant effect of vascular health risk on qT1 values (M1: χ2(1)=1.35, p=0.245; S1: χ2(1)=0.25, p=0.621) and no significant interaction effect between vascular health risk and distance to the nearest vein in older adults (M1: χ2(5)=5.22, p=0.389; S1: χ2(5)=1.70, p=0.889) (see **Figure 4** for individual values).

**Figure 4.**
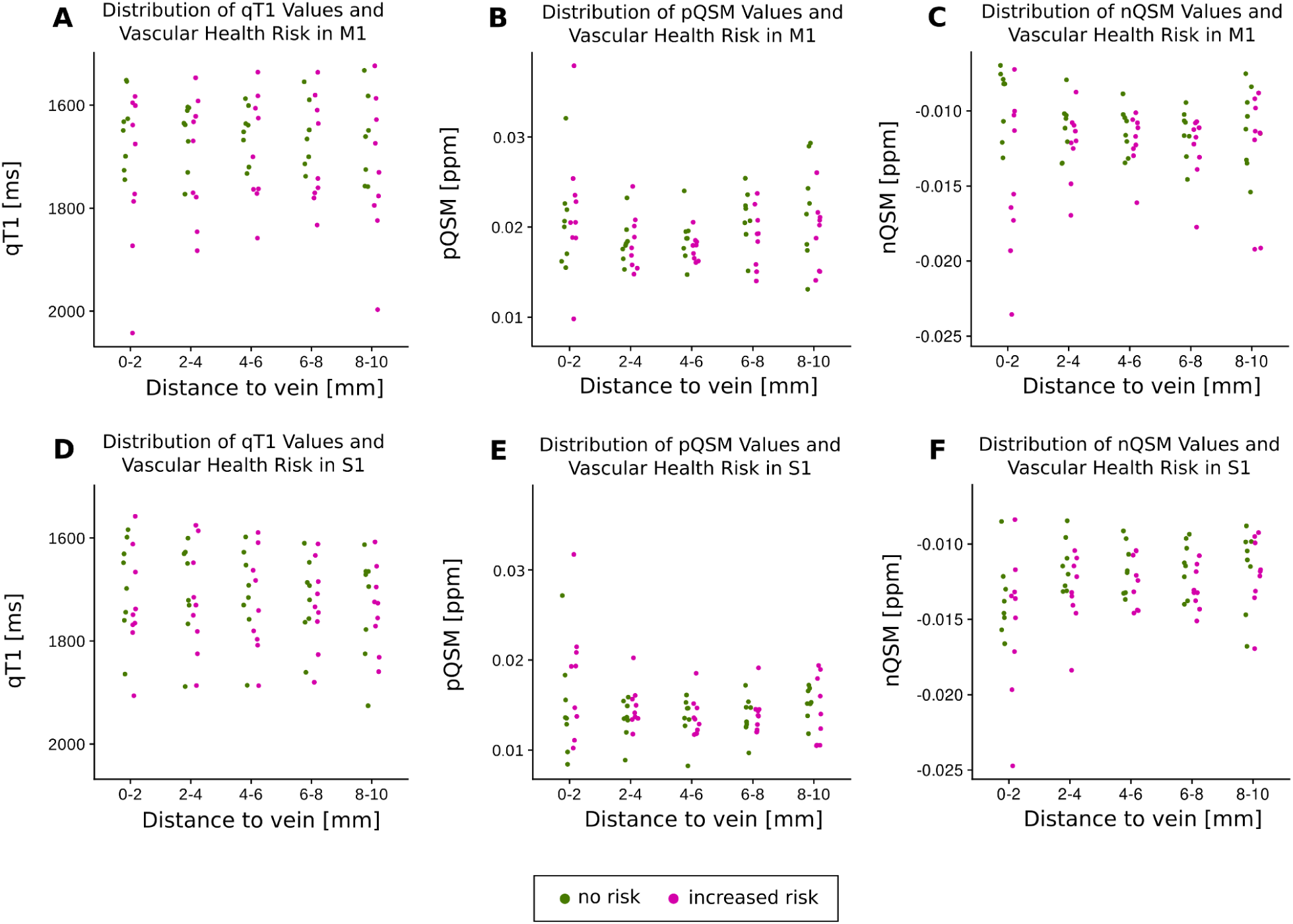
Distribution of vascular health risks in older adults. Individual vascular health risks (i.e., binary variable indicating the presence of hypertension and/or signs of cerebral small vessel diseases according to the STRIVE-2 criteria; green: no risk, magenta: increased risk) shown in relation to the distance to the nearest vein (given in millimetres). **A**: Distribution of vascular health risks for older adults (n=17) in primary motor cortex (M1) in relation to quantitative T1 values (qT1, given in milliseconds, i.e. ms). **B**: Same as in A but for positive QSM values (pQSM, given in parts per million, i.e. ppm). **C**: Same as in A but for negative QSM values (nQSM, given in ppm). **D**: Distribution of vascular health risks for older adults in primary somatosensory cortex (S1) in relation to qT1 values. **E**: Same as in D but for pQSM values. **F**: Same as in D but for nQSM values.

### Cortical Iron Shows a Relationship to the Distance to the Nearest Vein

To test for the main effect of venous distance on pQSM values and a possible interaction with age, we computed the same ANOVA as reported above on pQSM values. As expected, there is a significant main effect of age with higher pQSM values (more iron) in older compared to younger adults, a significant main effect of brain area with higher pQSM values in M1 compared to S1, and a significant main effect of cortical depth driven by reduced pQSM values in deeper cortical depths (see **Table 3**, **Figure 5**).

**Figure 5.**
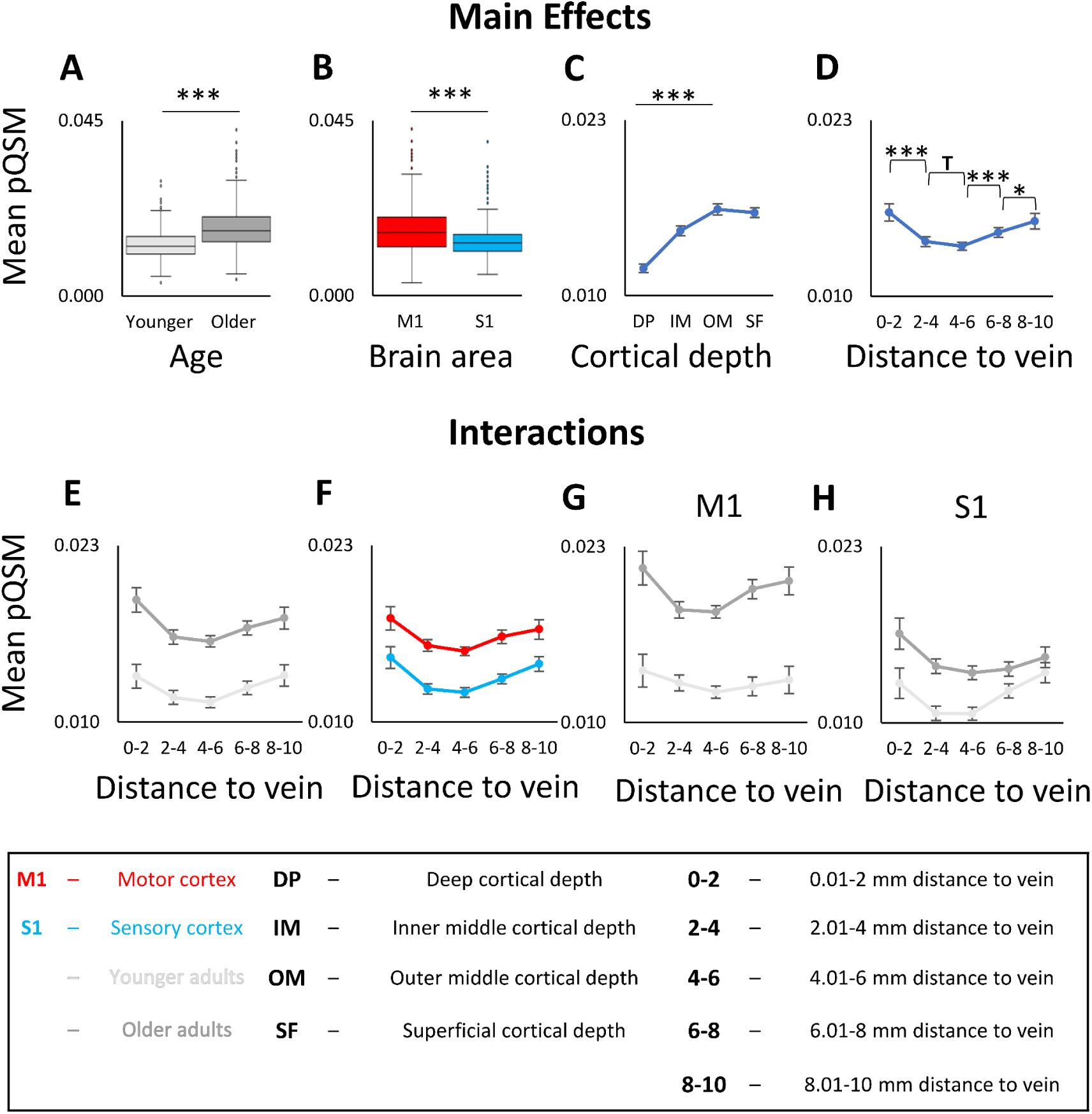
Interaction between age, brain region, cortical depth and venous distance for pQSM values. **A:** Significant main effect of age on pQSM values. Shown are medians, interquartile ranges, and lower and upper quartiles for younger (n=18, *light grey*) and older (n=17, *dark grey*) adults. Dots mark outliers. pQSM values are given in parts per million (higher values indicate higher iron content). **B:** Significant main effect of brain area (M1 (*red*), S1 (*blue*)) on pQSM values. **C:** Significant main effect of cortical depth on pQSM values. Dots indicate mean pQSM values for different cortical depths averaged across age groups, distances, and brain areas. Error bars indicate standard errors of the mean (SEM). **D:** Significant main effect of venous distance on pQSM values (0-2 = 0.01-2 mm, 2-4 = 2.01-4 mm, 4-6 = 4.01-6 mm, 6-8 = 6.01-8 mm, 8-10 = 8.01-10 mm). **E:** No significant interaction effect between venous distance and age on pQSM values. **F:** No significant interaction effect between venous distance and brain area on pQSM values. **G/H:** No significant interaction effect between venous distance, age and brain area (**G:** data for M1, **H:** data for S1). Significant results of post-hoc t-tests to follow-up mixed-effects ANOVA results (see **Table 3** for statistical results) are marked by asterisks: * p≤0.05, ** p≤0.005, *** p≤0.0005 (uncorrected). Trends above p=0.05 are marked by a T.

**Table 3.**
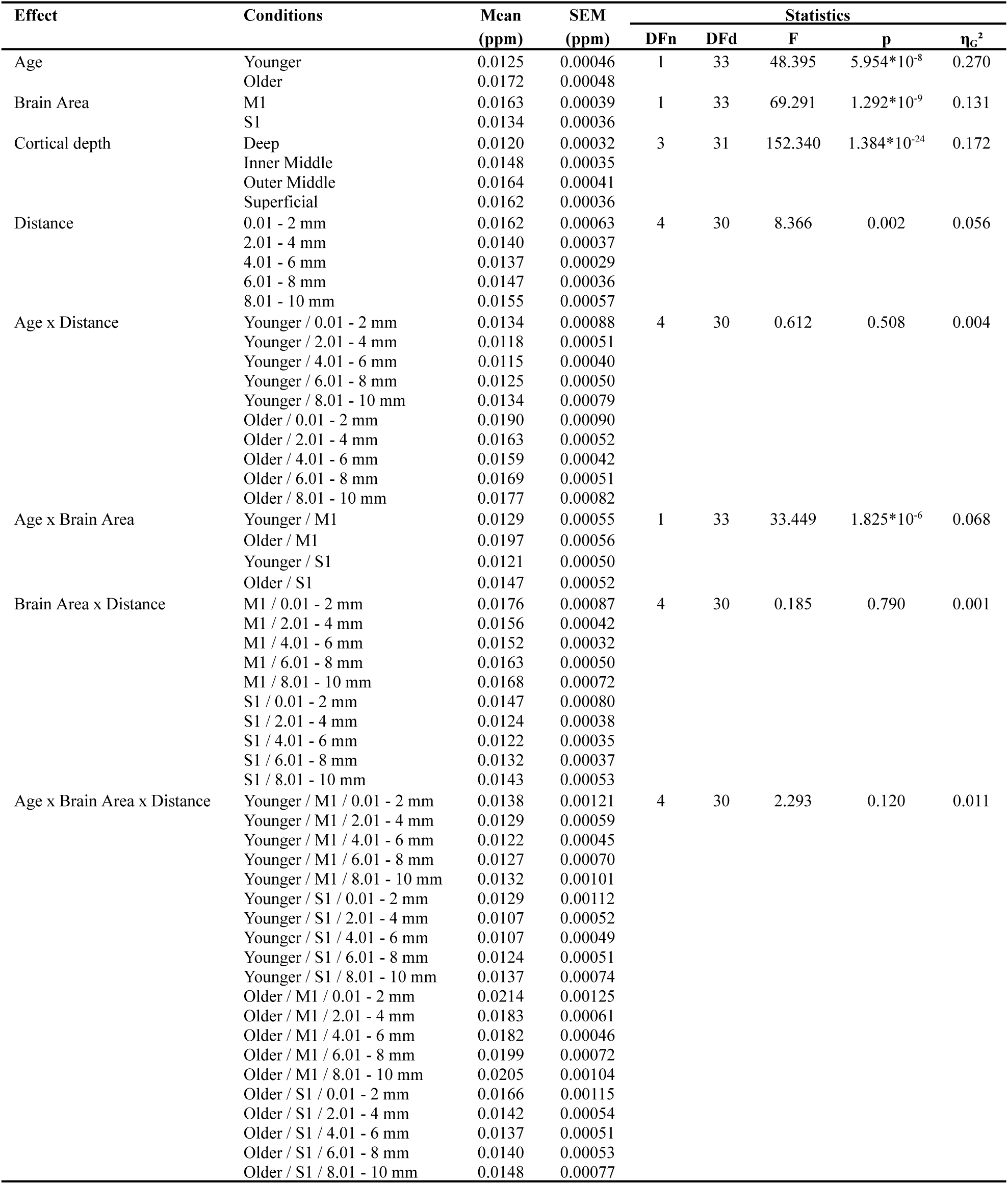
ANOVA Results for pQSM values. Results for the ANOVA on pQSM values with factors age (n=18 younger adults, n=17 older adults), cortical depth (SF, OM, IM, DP), brain area (S1, M1) and venous distance (5 distances). Given are mean pQSM values in parts per million (Mean in ppm) and standard error of the mean (SEM), degrees of freedom of the numerator (DFn), degrees of freedom of the denominator (DFd), p-values (p), and effect sizes (n_G_²). P-values ≤ 0.05 are considered significant.

There is also a significant main effect of venous distance (see **Table 3**, **Figure 5D**) driven by significantly higher pQSM values (more iron) in bin 0-2 compared to bin 2-4 (t(34)=4.388, p=1.052*10^-4^, p_Holm-Bonferroni_=4.208*10^-4^, d=0.73) and significantly lower pQSM values (less iron) in bin 4-6 compared to bin 6-8 (t(34)=-6.132, p=5.799*10^-7^, p_Holm-Bonferroni_=1.740*10^-6^, d=-1.01) and in bin 6-8 compared to bin 8-10 (t(34)=-2.289, p=0.028, p_Holm-Bonferroni_=0.056, d=-0.38). However, the latter result does not survive Holm-Bonferroni correction. In addition, there is a trend towards higher pQSM values in bin 2-4 compared to bin 4-6 (t(34)=1.715, p=0.095, d=0.28). This shows that in both younger and older adults, there is a U-shaped relationship between cortical iron content and venous distance. Other than for qT1 values, this effect is not significantly modulated by brain area (see **Table 3** and **Figure 5F**). The venous distance explains 5.6% of the variance in the cortical pQSM signal.

Other than for qT1 values, there is no significant interaction between age and venous distance (see **Table 3**). These results are confirmed by permutation t-tests (see **Supplementary Table 5**). Instead, age effects differ significantly between brain areas (see **Table 3**): There are more pronounced age effects in M1 compared to S1 (M1: younger adults minus older adults =-0.0067 ppm, t(33)=-8.860, p=4.505*10^-11^, p_Holm-Bonferroni_=9.01*10^-11^, d=-2.80; S1: younger adults minus older adults =-0.0026 ppm, t(33)=-3.416, p=0.008, p_Holm-Bonferroni_=0.008, d=-1.18; see also **Figure 3**).

These results are confirmed when removing one older participant with a MoCA score below the cut-off for healthy ageing (see **Supplementary Table 6**) and when excluding the closest distance (bin 0-2), except that for the latter there is a significant three-way interaction between age, brain area and venous distance (see **Supplementary Table 7**). This interaction is driven by the less pronounced U-shaped relationship between pQSM values and venous distance when the closest distance is removed. There are age-related differences in cortical iron content that depend on the distance to the nearest vein where this effect is different in M1 versus S1 (M1: more iron in older adults with increasing venous distances (older adults minus younger adults): 2.01-4 mm=0.0054 ppm, 4.01-6 mm=0.0059 ppm, 6.01-8 mm=0.0072 ppm, 8-10=0.0073 ppm; S1: more iron in older adults with decreasing venous distances (older adults minus younger adults): 2.01-4 mm=0.0035 ppm, 4.01-6 mm=0.0030 ppm, 6.01-8 mm=0.0016 ppm, 8.01-10 mm=0.0011 ppm). Taken together, age effects on iron content are stronger in M1 compared to S1. When excluding the closest distance, it becomes apparent that higher iron content in older adults is more pronounced at distances farther away from a vein in M1 and closer to a vein in S1.

Controlling for effects of vascular health, comparisons of random intercept models reveal no significant effect of vascular health risk on pQSM values (M1: χ2(1)=0.64, p=0.423; S1: χ2(1)=0.66, p=0.418) and no significant interaction effect between vascular health risk and distance to the nearest vein in older adults (M1: χ2(5)=4.43, p=0.489; S1: χ2(5)=5.13, p=0.400) (see also **Figure 4** for individual values).

Finally, we correlated pQSM with qT1 values for all area-by-distance conditions separately for younger and older adults (see **Supplementary Figure 3, Supplementary Table 8**). Particularly at distances close to the nearest vein (bin 0-2) the results indicate only weak correlations in both age groups.

### Cortical nQSM Values Show a Relationship to the Distance to the Nearest Vein

To test for the main effect of venous distance on nQSM values, and a possible interaction with age, we computed the same ANOVA as reported for qT1 and pQSM on nQSM values. As expected, there is a significant main effect of age with more negative nQSM values (higher calcium/protein content) in older adults compared to younger adults (see **Figure 3** for effect sizes). This is confirmed by permutation t-tests (see **Supplementary Table 9**). There is also a significant main effect of brain area with more negative nQSM values (higher calcium/protein content) in S1 compared to M1 (see **Supplementary Table 10**, **Supplementary Figure 4B**) and a significant main effect of cortical depth driven by more negative nQSM values in deep and superficial compared to middle cortical depth (inverted U-shape, see **Supplementary Figure 4C**).

There is also a significant interaction between brain area and venous distance on nQSM values (see **Supplementary Table 10**). This is driven by a significant effect of venous distance in S1, showing more negative nQSM values in bin 0-2 compared to bin 2-4 (t(34)=-4.196, p=1.841*10^-5^, p_Holm-Bonferroni_= 7.364*10^-5^, d=-0.69). All other comparisons are not significant or offer trends (more negative nQSM values in bin 6-8 compared to bin 8-10 (t(34)=-1.936, p=0.061, d=-0.32) in S1; less negative nQSM in bin 4-6 compared to bin 6-8 (t(34)=1.943, p=0.060, d=0.32) in M1). There is neither a significant interaction between venous distance and age nor between venous distance, age and brain area (see **Supplementary Table 10**). When excluding the closest distance (bin 0-2), there is still a significant main effect of cortical depth and an interaction between brain area and venous distance, but no main effect of brain area (see **Supplementary Table 11**).

Controlling for effects of vascular health, comparisons of random intercept models revealed no significant effect of vascular health risk on nQSM values (M1: χ2(1)=3.21, p=0.073; S1: χ2(1)=1.75, p=0.19). However, there was a significant interaction effect between vascular health risk and distance to the nearest vein on nQSM values in M1, but not in S1 in older adults (M1: χ2(5)=18.02, p=0.003; S1: χ2(5)=3.67, p=0.598), indicating that older adults with vascular health risk factors exhibit more negative nQSM values in M1 at distances close to the nearest vein (0.01-2 mm) compared to older adults without vascular health risk factors (see also **Figure 4** for individual values).

### Interaction between Brain Microstructure and Venous Distance is not restricted to Sensorimotor Cortex

To investigate whether the above reported effects also occur more widely throughout the frontal cortex or are restricted to the sensorimotor system, we conducted additional analyses incorporating the SFG, CMF and RMF. Please note that additional brain areas, for example in the temporal cortex, could not be integrated into these analyses given our slab acquisition did not cover the entire brain. We computed an ANOVA with the factors, age (younger adults, older adults), brain area (S1, M1, SFG, CMF, RMF), cortical depth (SF, OM, IM, DP) and venous distance (bin 0-2, bin 2-4, bin 4-6, bin 6-8, bin 8-10) on qT1 and pQSM values. We replicate the main effect of venous distance on qT1 values (see **Supplementary Figure 5, Supplementary Table 12**) and pQSM values (see **Supplementary Figure 6, Supplementary Table 13**), and the significant interaction between age and venous distance on qT1 values (see **Supplementary Figure 5, Supplementary Table 12**) when three additional brain areas were integrated into the analyses. Whereas similarities and differences between brain areas for each structural marker can be inspected in detail in **Supplementary Figures 5-7 and Supplementary Tables 12-14**, the replication of the main effects and interactions show that an influence of the venous architecture on brain microstructure is not restricted to the sensorimotor system.

## Discussion

Our data shows a dependence of standard measures of cortical microstructure on the distance to the nearest vein both in younger and older adults. In addition, with respect to ageing, we tested three hypotheses: Cortical locations that are more distant to veins are more prone to age-related degeneration, and cortical locations that are closer to a vein are more protected from age-related decline (H1), cortical locations that are closer to veins are more prone to age-related degeneration, and cortical locations that are farer away from a vein are more protected from age-related decline (Hypothesis 2, H2), finally, cortical degeneration occurs independently of the distance to the nearest vein (H3). Whereas H3 is supported for age-related differences in QSM values, an interaction between venous distance and age occurs for qT1 values. When interpreting low qT1 values as markers of high myelin^17^, and high qT1 values as markers of low myelin (hence neurodegeneration^59^), our data supports H1 for cortical myelin. Other interpretations of the lower qT1 values that older adults show compared to younger adults in voxels closer to veins are discussed below. Together, this study shows that the local venous architecture explains a significant amount of variance in standard measures of cortical microstructure and needs to be considered in neurobiological models of cortical ageing and microstructural brain organisation.

We report a significant interaction effect between age and venous distance on qT1 values. The interaction is driven by older adults showing lower qT1 values (interpreted as more myelin)^17^ in voxels closer to a vein compared to younger adults. In addition, in M1, older adults show higher qT1 values (interpreted as less myelin)^59^ in voxels farther away from a vein compared to younger adults. This pattern of results, and the interpretation of qT1 values representing intact cortical myelination, confirms H1 for cortical qT1 values. Prior studies show either reduced or higher cortical myelination in older compared to younger adults^9–11,70,71^ or no significant differences at all.^59^ Those contrary findings may relate to hidden variables that explain a significant amount of variance but have been disregarded so far. Our data indicate that the venous architecture, specifically the distance to the nearest vein, acts as such a hidden variable, and may explain prior contradictory findings on age-related changes in cortical T1 values.

Possible neuronal mechanisms are that increased metabolism close to blood vessels may prevent age-related myelin loss, and/or ageing blood vessels particularly reduce supply to locations that are farther away from major branches. Myelin itself can be a driver for increased metabolism, since it is a highly demanding structure.^72^ The venous architecture would in this view be a protective structure for older adults’ cortical microstructure, specifically with respect to myelin loss. Perosa et al.^3^ hypothesised that in the hippocampus, a steady arterial vascular supply constitutes a ‘vascular reserve’ mechanism in cases of pathology, relating to larger hippocampal volume^4^ and preserved cognitive abilities in patients with cerebral small vessel disease.^73^ Our results indicate that a similar vascular reserve mechanism can be assigned to veins.

Our study cannot clarify if the lower qT1 values in older adults in cortical locations close to veins indeed reflect higher myelination, and if this is an adaptive or maladaptive process, and/or a marker of vascular reserve, compensation or even pathology. Our finding that older adults with vascular health risk factors exhibit higher nQSM values in M1 at distances close to the nearest vein compared to older adults without vascular health risk factors indicates that the substance close to the vein may also contain markers of calcification and/or protein accumulation. To clarify whether increased substances close to a vein in older adults relate to better or worse cognitive performance and brain health, a larger cohort needs to undergo ultra-high resolution imaging in combination with assessments of cognitive functions and markers of neurodegeneration, such as amyloid and tau PET imaging.^74^ In case future research would identify the lower qT1 signal in older adults close to veins as maladaptive or as marker of neurodegeneration, H2 would be accepted for qT1. Additionally, investigating patients with exceptionally high or low myelin content close to vessels in a longitudinal study may allow addressing its effect on brain health.

Both younger and older adults show a U-shaped relationship between pQSM values (iron) and venous distances, driven by highest pQSM values at cortical voxels closest and farthest away from a vein. This effect is not significantly modulated by age, confirming H3. However, methodological limitations need to be considered: Whereas we here focus on veins at the mesoscopic scale, it is possible that microvessels located close to larger veins increase pQSM values. Increased pQSM values at distances close to a vein need to be treated with caution and could potentially be driven by partial volume artefacts. Therefore, we recalculated all analyses without the closest distance. When doing so, the cortical myelin effects as described above remain, but there is a significant interaction between age, brain area, and venous distance on pQSM values. This is driven by opposing relationships between age-related iron content and venous distance in M1 versus S1: Whereas age-related differences in iron content are larger at greater venous distances in M1, they are larger at smaller venous distances in S1. This would confirm H1 for M1 and H2 for S1, rather than confirming H3. Given the resolution was 0.5 mm isotropic, which limits the amount of small veins that can be classified, the present data cannot clarify the relationship between age-related iron accumulation and venous distance, but warrants further research using higher resolutions (0.35 mm or higher).^75^

When brain areas are inspected in isolation, differences become apparent. The U-shaped relationship between venous distance and pQSM values is present in the M1, S1, RMF and CMF, but is not present in the SFG. The interaction between venous distance and age on qT1 values is driven by the M1, S1 and CMF, but is not present in the SFG and RMF. The reason for these differences is at present unclear. Given the sensorimotor cortex shows higher myelination than the other three investigated cortical areas, effects of venous distances may be easier to detect in the sensorimotor cortex. However, given the CMF shows an interaction between age and cortical distance, this cannot be the only explanatory factor. Alternatively, the effect of venous architecture might be particularly strong in areas with pronounced age effects, which are almost absent in the RMF and weak in the SFG. In addition, in the SFG, the effect of venous distance on pQSM values shows an opposing direction far away from veins to the other investigated areas in younger adults, which may be explained by differences in vascular plasticity^76^ or connectivity.^77^ Whereas our results therefore show that the reported effects are not restricted to the sensorimotor cortex, future research should clarify the factors that contribute to areal differences presented in this study.

With respect to further study limitations, the participant number was relatively low but motivated by previous layer-dependent 7T-MRI studies using quantitative in-vivo proxies to describe the microstructural cortex architecture^16,49^, and is above previously reported sample sizes. To reduce type I and type II errors, we applied corrections for multiple comparisons and reported statistical trends. In addition, we conducted permutation tests, known to be superior to conventional tests when investigating small samples^78,79^, and we reported effect sizes. 7T-MRI has superior statistical power compared to MRI at lower field strengths, showing effects at the single participant level^80,81^ (see **Supplementary Figures 1, 2**). Nevertheless, the inclusion of more participants would have benefitted the study with respect to generalizability and robustness. Moreover, the image resolution of 0.5 mm isotropic does not allow to identify the entire venous network. For a comprehensive investigation of the neuronal mechanisms that underlie the interaction between blood vessels and cortical microstructure, also the arterial blood supply patterns need to be investigated, and should be complemented by blood flow assessments. Analyses of time-of-flight MRI measurements would benefit future studies. Moreover, previous studies pointed out that hypertension and alterations in brain vasculature relate to myelin loss and M1 pathology^5,82–85^, and 9/17 older adults show global signs of vascular health risk. However, apart from nQSM values at the closest distance in M1, these indicators do not significantly predict S1 and M1 microstructure in older adults, and may therefore not account for the main findings. Assessing the potential impact of hypertension and other vascular disorders on cortical microstructure is an important topic for future studies.

Taken together, we show that the venous architecture interacts with cortical microstructure and age-related changes thereof. This is an important insight as most studies do not analyse cortical alterations as a function of venous or arterial distance, which may lead to misinterpretations of present (or absent) effects. Our study provides evidence for the importance of the human venous architecture in understanding cortical function in health and disease.

## Data availability

MRI data available upon request, requirements are a formal data sharing agreement and need to submit a formal project outline.

## Code availability

Code for small vessel segmentation hysteresis thresholding: https://gitlab.com/hmattern/omelette

## Author contributions

**Christoph Knoll:** Formal analysis, Investigation, Writing Original Draft -review & editing **Juliane Doehler:** Data collection, Formal analysis, Methodology, Investigation, Writing -review & editing **Alicia Northall:** Formal analysis, Writing -review & editing **Stefanie Schreiber:** Conceptualization, Funding acquisition, Writing -review & editing **Johanna Rotta:** Formal analysis **Hendrik Mattern:** Methodology, Software, Formal analysis, Writing -review & editing **Esther Kuehn:** Conceptualization, Investigation, Funding acquisition, Supervision, Writing -review & editing.

