## Supplemental Methods, Tables and Figures for "Age-Related Differences in Human Cortical Microstructure Depend on the Distance to the Nearest Vein"

### Supplementary Material

#### Supplementary Methods

Tactile detection task

Tactile detection thresholds of the right index fingertip were assessed using fine hair stimuli (Semmes-Weinstein monofilaments; Baseline, Fabrication Enterprises Inc., White Plains, NY, USA; stimuli intensities: 0.008 g, 0.02 g, 0.04 g, 0.07 g, 0.16 g, 0.4 g, 0.6 g, 1.0 g, 1.4 g, 2.0 g). Stimuli were manually applied to a predefined skin area (circle with 2mm diameter) at an angle of approximately 90 degree, for one second.[1] Stimulus application was guided by auditory cues (Psychophysics Toolbox for MATLAB R2017b).[2] All participants sat in front of a screen that signalled the beginning and ending of stimulus intervals and listened to white noise via headphones. The right hand (palm facing upwards) was fixated on a pillow behind a paper wall, preventing participants from seeing their own hand and the experimenter. In a two-alternative forced choice paradigm, participants chose one of two possible time intervals that contained the stimulation (randomly applied in the first or second interval).[3,4] Application of tactile monofilaments followed a 3-down/1-up staircase with two interleaved staircases, one starting at a 0.02 gram and the other starting at 0.4 gram. The stimulus weight was increased by one step after each error, and decreased by one step after every three correct responses, until stable performance was reached.[4] The experiment was finished when for the last 30 trials the standard variation in stimulus intensity was 1 step or less[4], or when the maximum number of 100 trials was reached. The participant‘s tactile detection threshold was defined as the mean stimulus intensity across reversal points (change of response from correct to incorrect or incorrect to correct) within the period of stable performance (i.e., the last 10 trials).[3] The experiment took approximately 12 min. Stimulus intensities were transformed logarithmically on a 1/10th milligram scale (log 10 0.1mg). Lower values indicate higher tactile sensitivity to mechanical forces.

Two-point discrimination task

We also assessed tactile 2-point discrimination performance of the right index fingertip.[5–7] Stimulation was applied by two rounded pins (diameter=0.4 mm) simultaneously touching the skin. A custom-made, fully automatic stimulation device moved the pins up and down (controlled by Presentation version 16.5, Neurobehavioral Systems, Inc., Albany, CA, USA). The amplitude of pin movement was adjusted to the individual detection threshold, but was at least set to 1.2 mm. Spacing between pins ranged from 0.7 to 2.8 mm (in steps of 0.3 mm) for younger adults and from 0.7 to 6.3 mm (in steps of 0.8 mm) for older adults. A single pin was included as control condition. Pin spacing was vertically adjusted by a rotating disc containing all possible conditions (n=9). In a two-alternative forced-choice paradigm, pin conditions were pseudo-randomly presented. Participants indicated whether they perceived one or two single pins touching their fingertip. They were instructed to give the answer 'two pins felt' only if they were certain. The hand was covered by a white box during the task. Each run included 90 trials (10 repetitions per pin condition). Unique sequences of pin spacing conditions were used per participant and run. All participants completed two runs. Intertrial intervals were pseudo-randomized and varied between 1 to 5 seconds. 2PD thresholds were calculated based on the second run. Answers "two pins felt" were fitted as percentages across ascending pin distances. A binary logistic regression was used to fit the data (glmfit function, Statistics Toolbox for MATLAB R2017b). The 2PD threshold was taken from the pin distance where the 50 percent level crossed the fitted sigmoid curve.[5,6,8,9] Lower values indicate higher spatial acuity.

Sensorimotor integration

Sensorimotor integration was assessed with a custom made pressure sensor (held between the thumb and index finger of the right hand).[10] Reference forces that were to be matched ranged from 5% to 25% of the individual maximum grip force to avoid muscle fatigue.[11] Participants solved a visuo-motor matching task, demanding them to continuously adjust the grip force.[10,12] Applied forces were sampled at a frequency of 100 Hz and projected on screen at a refresh rate of 60 Hz. The task was controlled by the software package Presentation (version 16.5, Neurobehavioral Systems, Inc., Albany, CA, USA). Each trial contained a unique pseudorandomized sequence of 10 position changes at five different amplitudes (5%, 10%, 15%, 20%, 25% of maximum grip force), leading to a mean frequency of 0.25 Hz. After a period of task familiarisation[11,12], all participants performed the task for a total duration of 20 seconds. One run contained 15 trials with intertrial intervals of 10 seconds, lasting about 8 minutes in total. All participants performed two runs that were separated by a 5-minute resting period. After each trial, participants received feedback about their individual performance level on screen. We monitored the time (in seconds) the controllable bar was within a given percentage above (2.5%) and below (2.5%) the target line (upper edge of the reference bar)[10,11]. Higher values reflect better sensorimotor integration.

#### Supplementary Figures

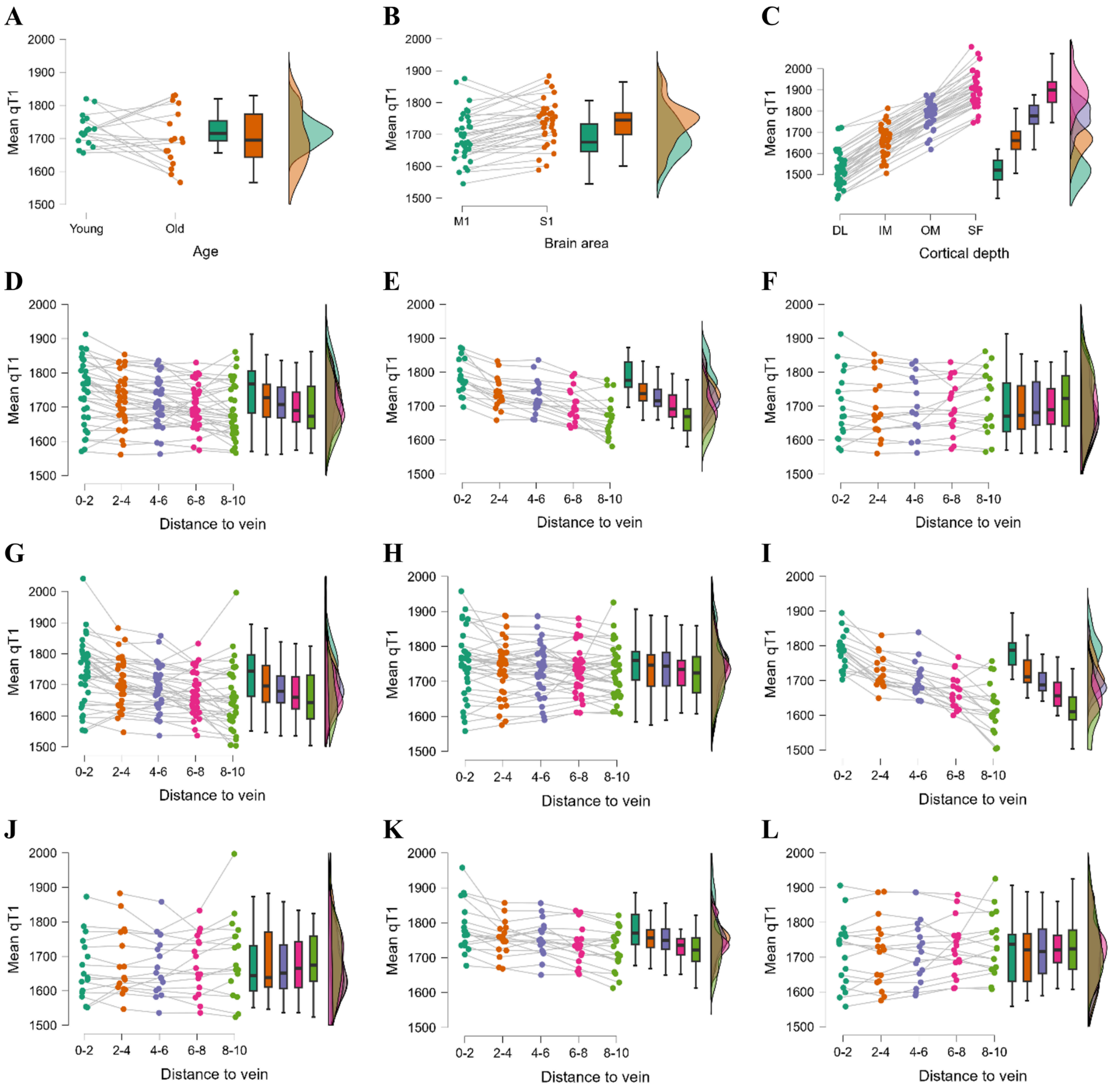


**Supplementary Figure 1. Descriptive raincloud plots of individual qT1 values for tested ANOVA conditions. A:** Younger versus older adults. **B:** M1 versus S1. **C:** Cortical depths. **D:** Distance to the nearest vein. **E:** Distance to the nearest vein for younger adults. **F:** Distance to the nearest vein for older adults. **G:** Distance to the nearest vein for M1. **H:** Distance to the nearest vein for S1. **I:** Distance to the nearest vein for younger adults in M1. **J:** Distance to the nearest vein for older adults in M1. **K:** Distance to the nearest vein for younger adults in S1. **L:** Distance to the nearest vein for older adults in S1. DL = deep cortical depth, IM = inner middle cortical depth, OM = outer middle cortical depth, SF = superficial cortical depth. M1 = primary motor cortex, S1 = primary somatosensory cortex. qT1 values given in milliseconds. Distance to the nearest vein given in millimetres.

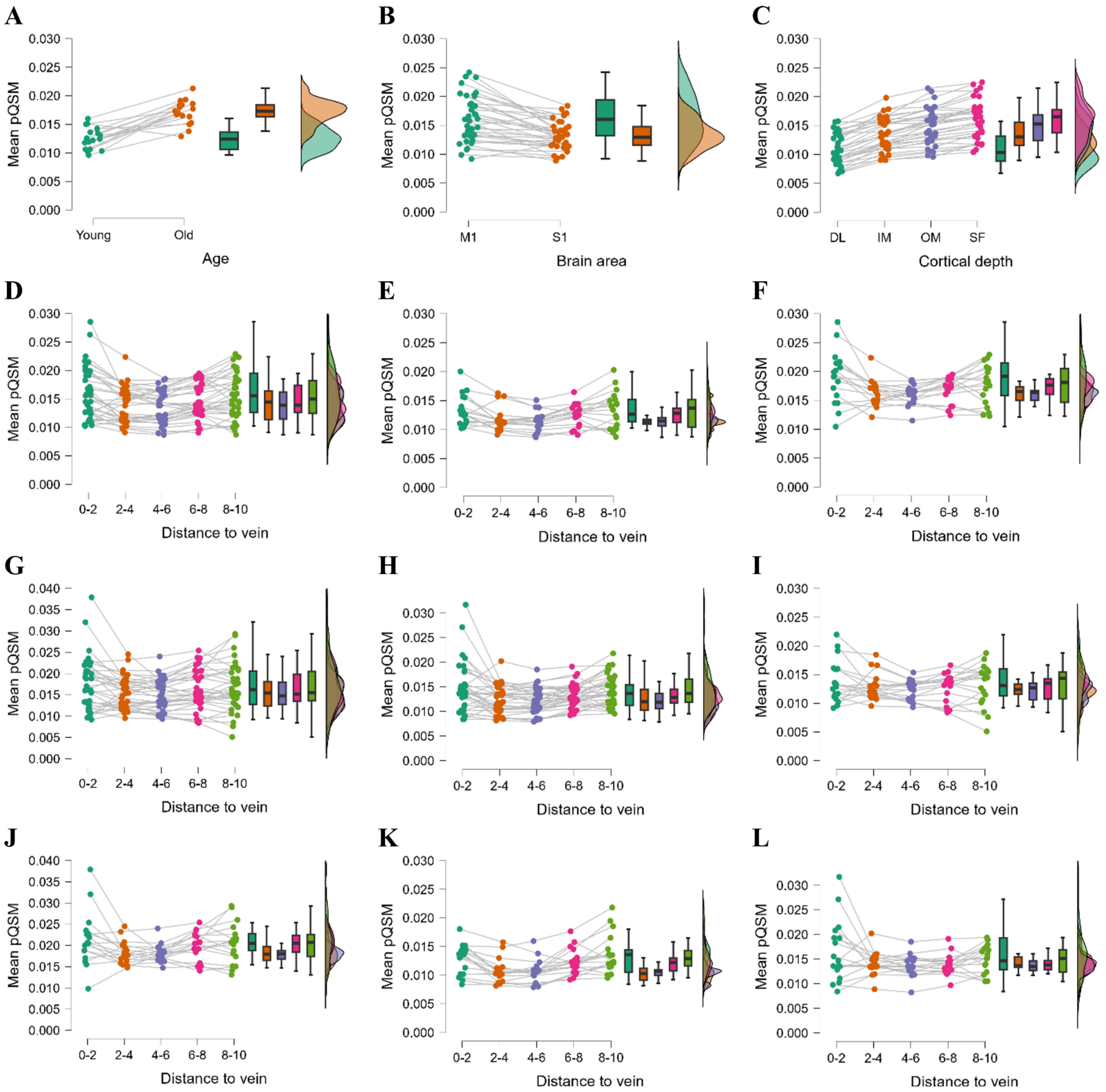


**Supplementary Figure 2. Descriptive raincloud plots of individual pQSM values for tested ANOVA conditions. A:** Younger versus older adults. **B:** M1 versus S1. **C:** Cortical depths. **D:** Distance to the nearest vein. **E:** Distance to the nearest vein for younger adults. **F:** Distance to the nearest vein for older adults. **G:** Distance to the nearest vein for M1. **H:** Distance to the nearest vein for S1. **I:** Distance to the nearest vein for younger adults in M1. **J:** Distance to the nearest vein for older adults in M1. **K:** Distance to the nearest vein for younger adults in S1. **L:** Distance to the nearest vein for older adults in S1. DL = deep cortical depth, IM = inner middle cortical depth, OM = outer middle cortical depth, SF = superficial cortical depth. M1 = primary motor cortex, S1 = primary somatosensory cortex. pQSM values given in parts per million. Distance to the nearest vein given in millimetres.

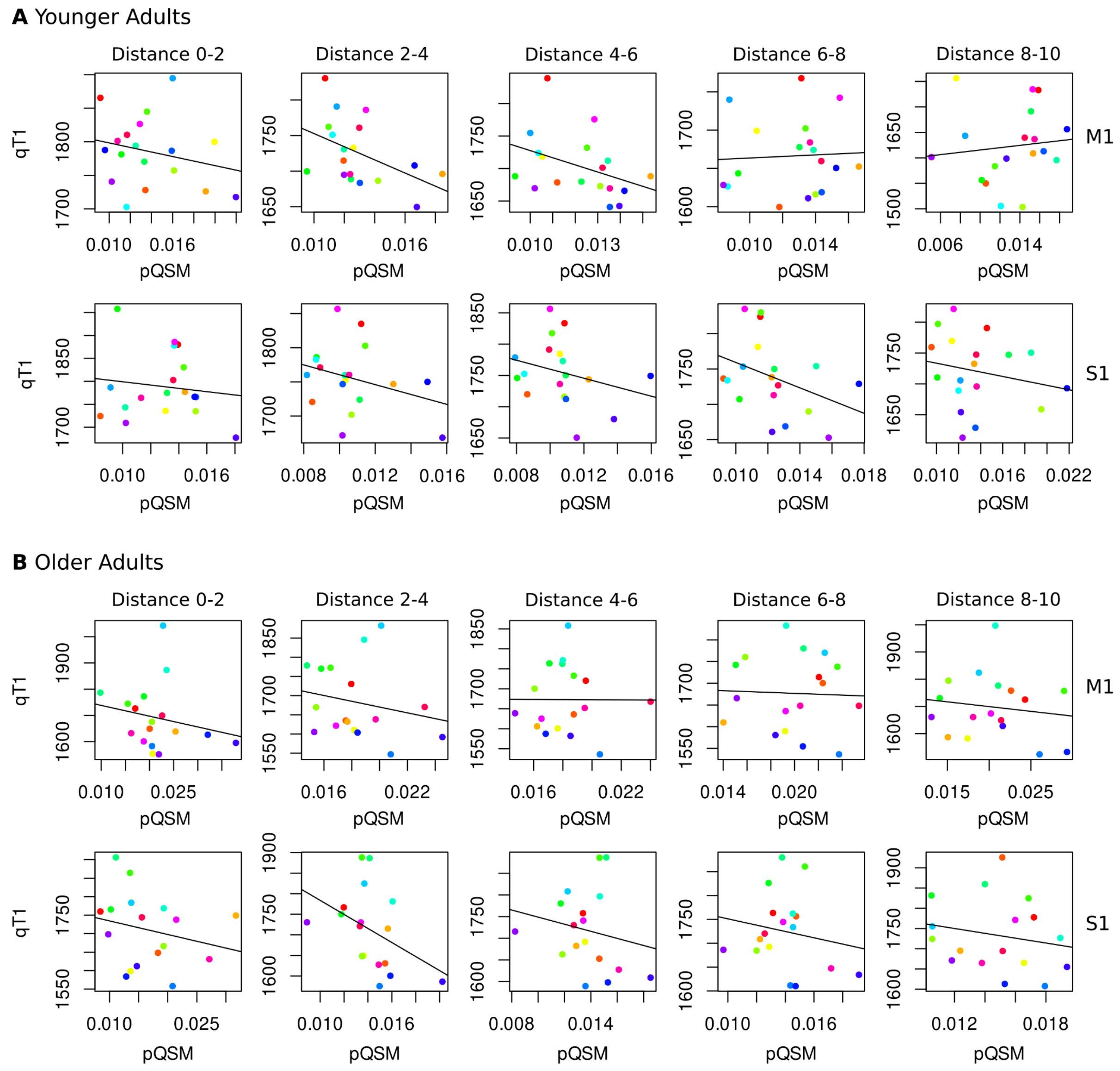


**Supplementary Figure 3. Relationship between qT1 and pQSM values.** Descriptive scatterplots showing individual qT1 values (milliseconds, y-axis) and pQSM values (parts per million, x-axis) for each brain area (M1, S1) by distance condition (0-2 mm, 2-4 mm, 4-6 mm, 6-8 mm, 8-10 mm). Different colours indicate different participants. Regression lines were generated based on simple linear regression models (qT1 ~ pQSM). For exact correlation coefficients see **Supplementary Table 8**. **A:** Scatterplots for younger adults (n=18). **B:** Scatterplots for older adults (n=17).

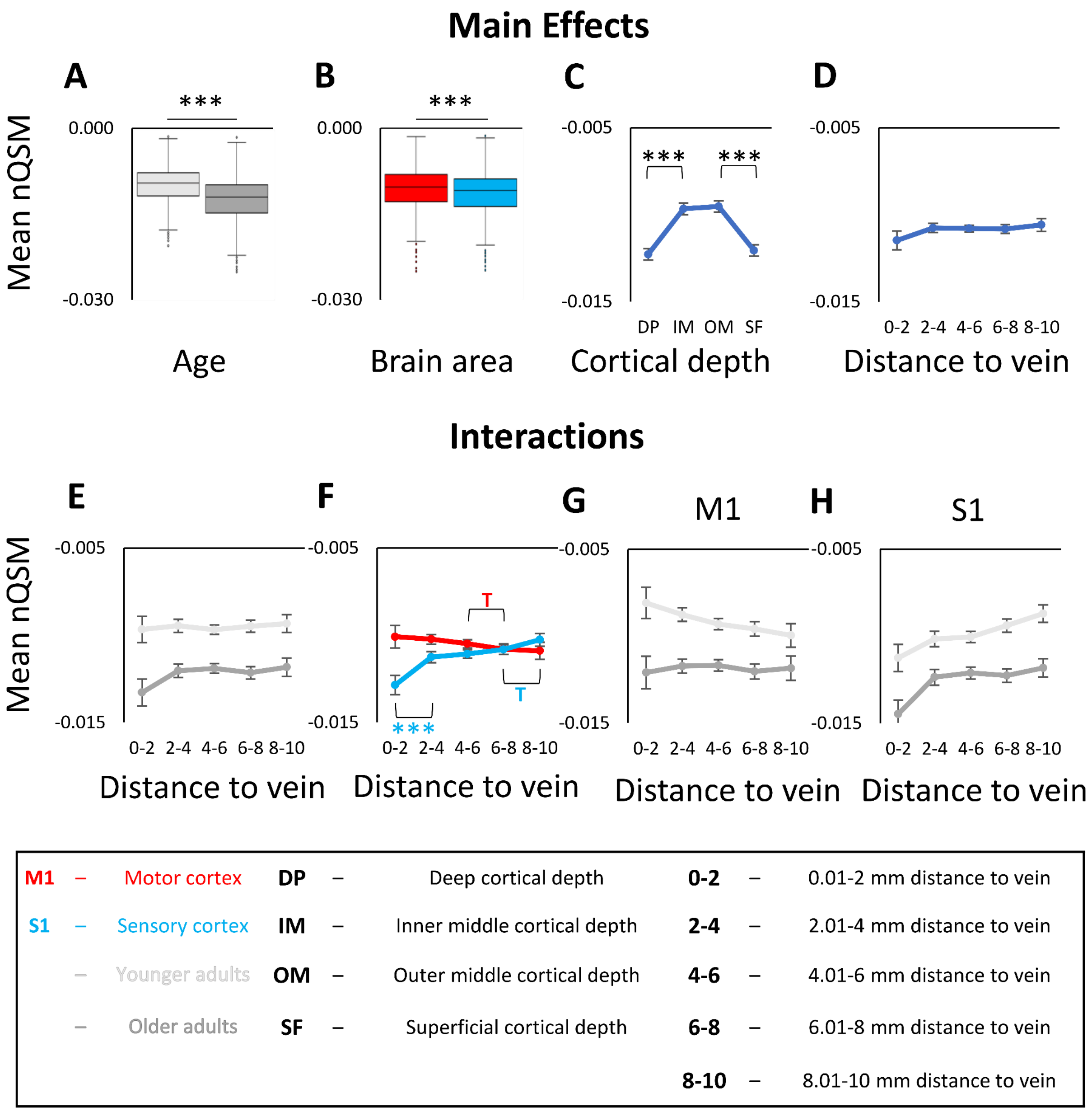


**Supplementary Figure 4. ANOVA results for nQSM values. A:** Significant main effect of age on nQSM values. Shown are medians, interquartile ranges, and lower and upper quartiles for younger (n=18, *light grey*) and older (n=17, *dark grey*) adults. Dots mark outliers. nQSM values are given in parts per million (lower values indicate higher diamagnetic contrast). **B:** Significant main effect of brain area (M1 (*red*), S1 (*blue*)) on nQSM values. **C:** Significant main effect of cortical depth on nQSM values. Dots indicate mean nQSM values for different cortical depths averaged across age groups, distances, and brain areas. Error bars indicate standard errors of the mean (SEM). **D:** No significant main effect of venous distance on nQSM values (0-2 = 0.01-2 mm, 2-4 = 2.01-4 mm, 4-6 = 4.01-6 mm, 6-8 = 6.01-8 mm, 8-10 = 8.01-10 mm). **E:** No significant interaction effect between venous distance and age on nQSM values. **F:** Significant interaction effect between venous distance and brain area on nQSM values. **G/H:** No significant interaction effect between venous distance, age and brain area (**G:** data for M1, **H:** data for S1). Significant results of post-hoc t-tests to follow-up mixed-effects ANOVA results (see **Supplementary Table 10** for statistical results) are marked by asterisks: * $p \leq 0.05$, ** $p \leq 0.005$, *** $p \leq 0.0005$ (uncorrected). Trends above $p=0.05$ are marked by a T.

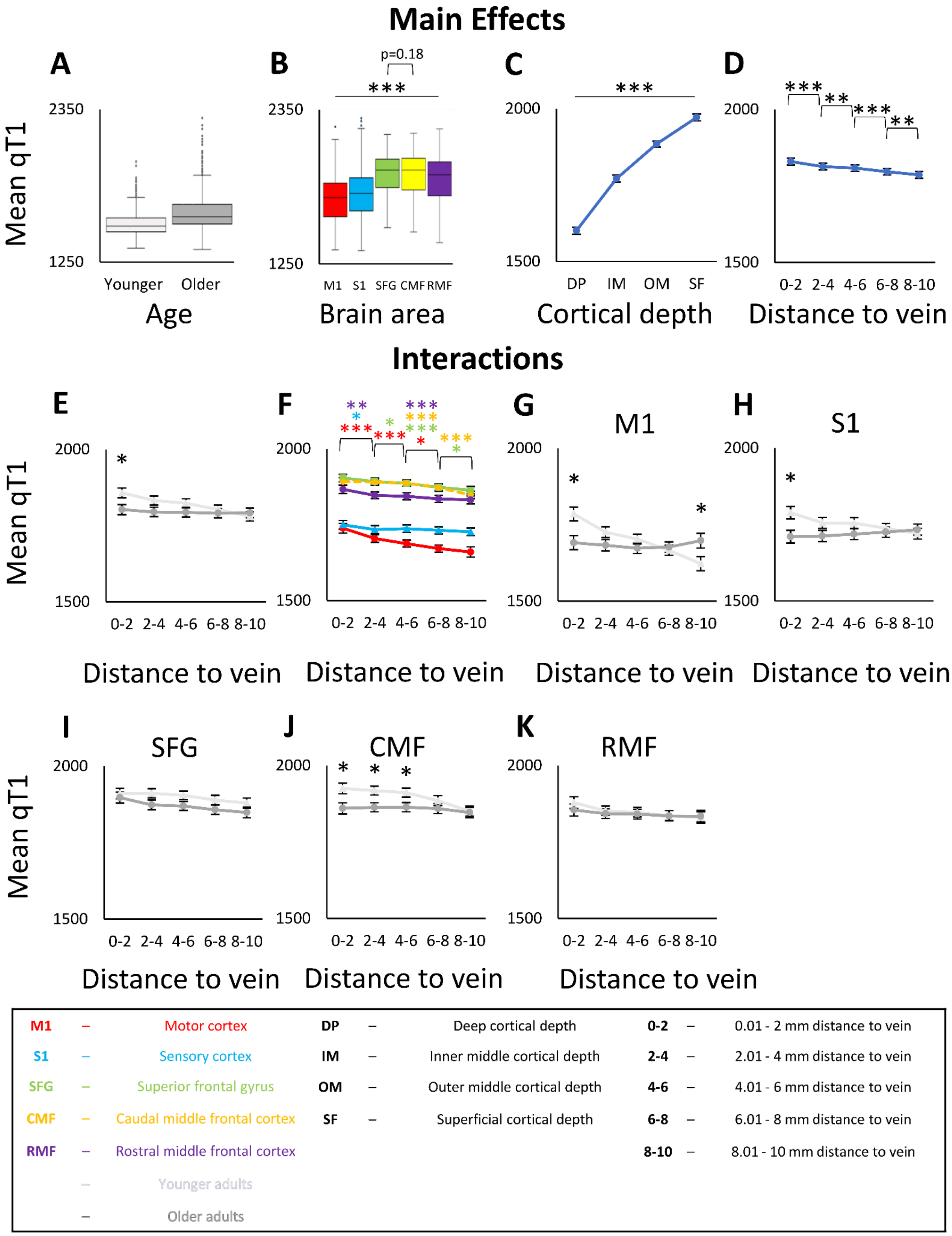


**Supplementary Figure 5. ANOVA results for qT1 values including primary motor cortex (M1), primary somatosensory cortex (S1), superior frontal gyrus (SFG), caudal middle frontal cortex (CMF) and rostral middle frontal cortex (RMF).** qT1 values are given in milliseconds (lower values indicate higher myelin content). **A:** No significant main effect of age on qT1 values. Shown are medians, interquartile ranges, and lower and upper quartiles for younger (n=18, *light grey*) and older (n=17, *dark grey*) adults. Dots above a box mark outliers. **B:** Significant main effect of brain area (i.e., M1 (*red*), S1 (*blue*), SFG (*green*), CMF (*yellow*), RMF (*violet*)) on qT1 values. **C:** Significant main effect of cortical depth on qT1 values; dots represent mean qT1 values for the different cortical depths averaged across age groups, distances, and brain areas. Error bars show standard errors of the mean (SEM). **D:** Significant main effect of venous distance on qT1 values (0-2 = 0.01-2 mm, 2-4 = 2.01-4 mm, 4-6 = 4.01-6 mm, 6-8 = 6.01-8 mm, 8-10 = 8.01-10 mm). **E:** Significant interaction effect between venous distance and age on qT1 values. **F:** Significant interaction effect between venous distance and brain area on qT1 values. **G-K:** Significant interaction effect between venous distance, age and brain area (**G**: data for M1, **H**: data for S1, **I**: data for SFG, **J**: data for CMF, **K**: data for RMF). Significant results of post-hoc t-tests to follow-up mixed-effects ANOVA results (see **Supplementary Table 12** for statistical results) are marked by asterisks: * $p \leq 0.05$, ** $p \leq 0.005$, *** $p \leq 0.0005$ (uncorrected). Trends above $p=0.05$ are marked by a T.

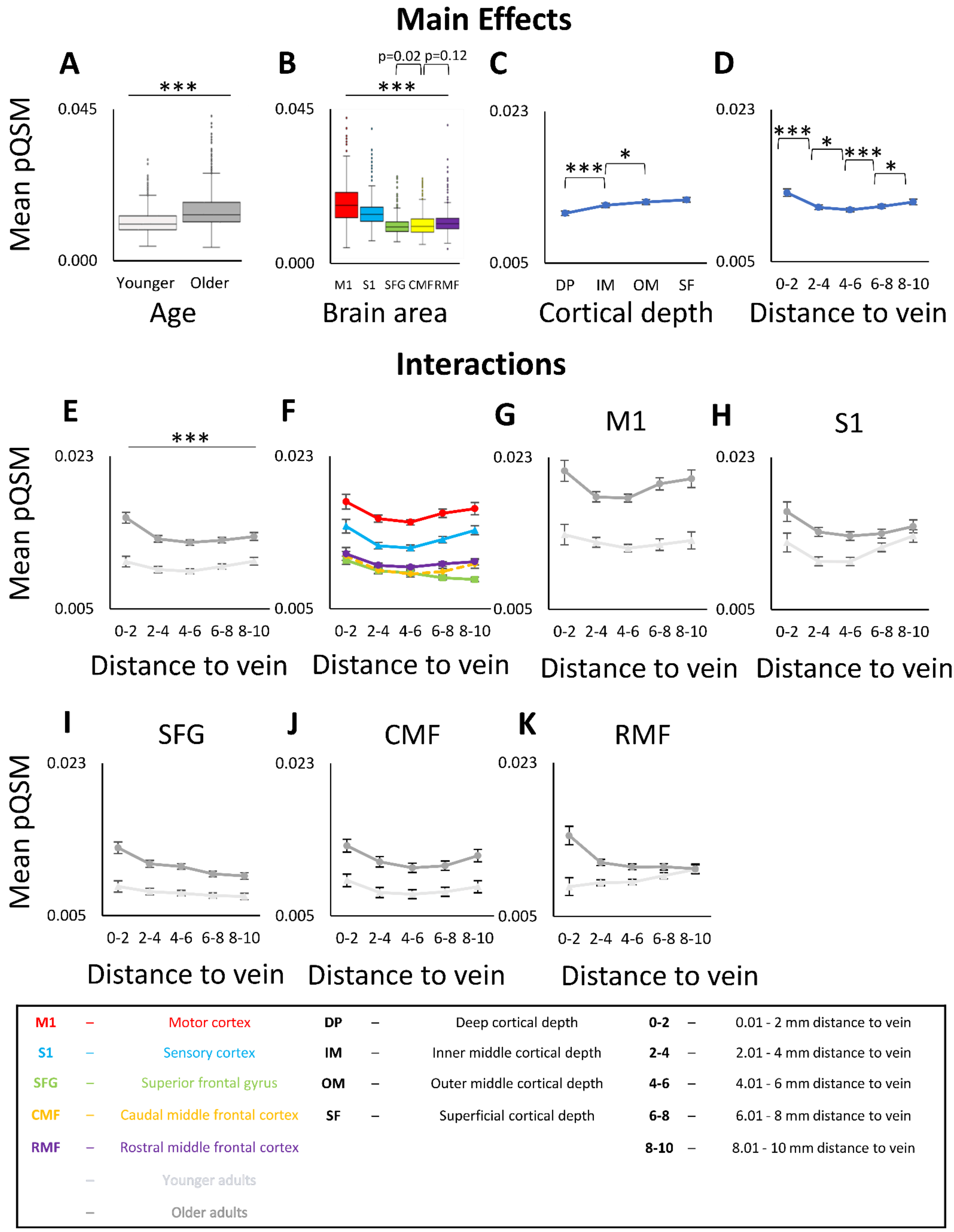


**Supplementary Figure 6. ANOVA results for pQSM values including primary motor cortex (M1), primary somatosensory cortex (S1), superior frontal gyrus (SFG), caudal middle frontal cortex (CMF) and rostral middle frontal cortex (RMF). A:** Significant main effect of age on pQSM values. Shown are medians, interquartile ranges, and lower and upper quartiles for younger (n=18, *light grey*) and older (n=17, *dark grey*) adults. Dots mark outliers. pQSM values are given in parts per million (higher values indicate higher iron content). **B:** Significant main effect of brain area (M1 (*red*), S1 (*blue*), SFG (*green*), CMF (*yellow*), RMF (*violet*)) on pQSM values. No significant effect for SFG-CMF and CMF-RMF comparisons. **C:** Significant main effect of cortical depth on pQSM values. Dots indicate mean pQSM values for different cortical depths averaged across age groups, distances, and brain areas. Error bars indicate standard errors of the mean (SEM). **D:** Significant main effect of venous distance on pQSM values (0-2 = 0.01- 2 mm, 2-4 = 2.01-4 mm, 4-6 = 4.01-6 mm, 6-8 = 6.01-8 mm, 8-10 = 8.01-10 mm). **E:** Significant interaction effect between venous distance and age on pQSM values. **F:** No Significant interaction effect between venous distance and brain area on pQSM values. **G-K:** No significant interaction effect between venous distance, age and brain area (**G:** data for M1, **H:** data for S1, **I**: data for SFG, **J**: data for CMF, **K**: data for RMF). Significant results of post-hoc t-tests to follow-up mixed-effects ANOVA results (see **Supplementary Table 13** for statistical results) are marked by asterisks: * $p \leq 0.05$, ** $p \leq 0.005$, *** $p \leq 0.0005$ (uncorrected). Trends above $p=0.05$ are marked by a T.

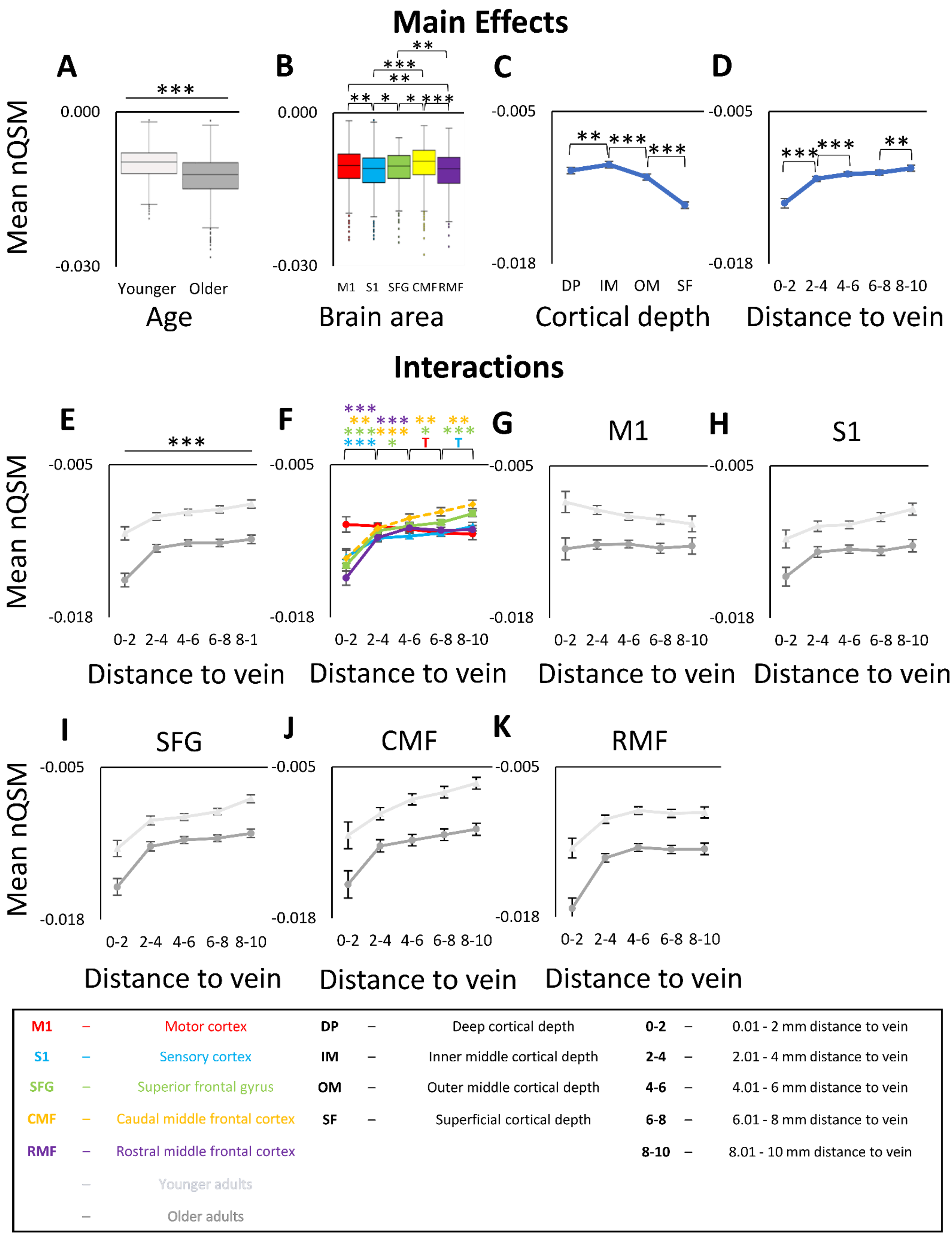


**Supplementary Figure 7. ANOVA results for nQSM values including primary motor cortex (M1), primary somatosensory cortex (S1), superior frontal gyrus (SFG), caudal middle frontal cortex (CMF), rostral middle frontal cortex (RMF). A:** Significant main effect of age on nQSM values. Shown are medians, interquartile ranges, and lower and upper quartiles for younger (n=18, *light grey*) and older (n=17, *dark grey*) adults. Dots mark outliers. nQSM values are given in parts per million (lower values indicate higher diamagnetic contrast). **B:** Significant main effect of brain area (M1 (*red*) versus S1 (*blue*), SFG (*green*), CMF (*yellow*), RMF (*violet*)) on nQSM values. **C:** Significant main effect of cortical depth on nQSM values. Dots indicate mean nQSM values for different cortical depths averaged across age groups, distances, and brain areas. Error bars indicate standard errors of the mean (SEM). **D:** Significant main effect of venous distance on nQSM values (0-2 = 0.01- 2 mm, 2-4 = 2.01-4 mm, 4-6 = 4.01-6 mm, 6-8 = 6.01-8 mm, 8-10 = 8.01-10 mm). **E:** Significant interaction effect between venous distance and age on nQSM values. **F:** Significant interaction effect between venous distance and brain area on nQSM values. **G-K:** No significant interaction effect between venous distance, age and brain area (**G**: data for M1, **H**: data for S1, **I**: SFG, **J**: data for CMF, **K**: data for RMF). Significant results of post-hoc t-tests to follow-up mixed-effects ANOVA results (see **Supplementary Table 14** for statistical results) are marked by asterisks: * $p \leq 0.05$, ** $p \leq 0.005$, *** $p \leq 0.0005$ (uncorrected). Trends above $p=0.05$ are marked by a T.

#### Supplementary Tables

| Task | Younger Adults | | | Older Adults | | | |
|---|---|---|---|---|---|---|---|
| | Mean ± SD | Value Range | # Extreme Outliers | Mean ± SD | Value Range | P2 (low MoCA) | # Extreme Outliers |
| Tactile Detection | 0.32 ± 0.06 | 0.23 - 0.44 | 0 | 0.27 ± 0.04 | 0.21 - 0.34 | 0.22 | 0 |
| Tactile 2-Point Discrimination | 0.56 ± 0.12 | 0.42 - 0.79 | 0 | 0.33 ± 0.07 | 0.19 - 0.51 | 0.19 | 0 |
| Sensorimotor Integration | 8.63 ± 2.27 | 4.83 - 12.11 | 0 | 5.78 ± 1.20 | 3.45 - 7.37 | 7.37 | 0 |

**Supplementary Table 1. Sensorimotor performance of younger and older adults.** Tactile detection (given in 1/[log10 0.1 mg]) and tactile two-point discrimination (given in 1/mm) were tested at the right index finger. Sensorimotor integration (given as precision grip accuracy in seconds) involved the thumb and the index finger. To ensure that higher values indicate better performance, tactile detection thresholds and tactile 2-point discrimination thresholds were reversed. Extreme outliers are defined as data points smaller or greater than the mean ± 3 standard deviations (SD). For the participant with the low MocA score (P2) individual values are given for the performed tasks.

| condition | diff ± se | p | 95% CI |
|---|---|---|---|
| qT1 M1_0_2 | 94.06 ± 35.72 | 0.007 | 24.88, 163.22 |
| qT1 M1_2_4 | 42.87 ± 26.03 | 0.105 | -8.01, 93.46 |
| qT1 M1_4_6 | 29.88 ± 23.60 | 0.217 | -16.30, 75.81 |
| qT1 M1_6_8 | -10.70 ± 23.33 | 0.660 | -56.15, 34.73 |
| qT1 M1_8_10 | -75.15 ± 34.96 | 0.029 | -143.00, -8.26 |
| qT1 S1_0_2 | 78.28 ± 31.55 | 0.012 | 16.94, 139.73 |
| qT1 S1_2_4 | 42.74 ± 26.57 | 0.111 | -8.60, 94.60 |
| qT1 S1_4_6 | 36. 45 ± 25.55 | 0.161 | -13.23, 86.24 |
| qT1 S1_6_8 | 13.15 ± 22.71 | 0.578 | -30.78, 57.31 |
| qT1 S1_8_10 | -12.73 ± 25.08 | 0.618 | -61.78, 36.23 |

**Supplementary Table 2. Results of permutation Welch two sample t-tests comparing qT1 values between younger adults (n=18) and older adults (n=17).** Results are given as permutation mean difference ± standard error (diff ± se) in milliseconds, permutation p-value (p) and 95% permutation confidence interval (95% CI).

| Effect | Conditions | Mean (ms) | SEM (ms) | Statistics | | | | |
|---|---|---|---|---|---|---|---|---|
| | | | | DFn | DFd | F | p | $\eta_G^2$ |
| Age | Younger | 1710.74 | 15.66 | 1 | 33 | 0.137 | 0.713 | 0.003 |
| | Older | 1702.41 | 16.12 | | | | | |
| Brain Area | M1 | 1680.95 | 12.30 | 1 | 33 | 41.597 | $2.585*10^{-7}$ | 0.086 |
| | S1 | 1732.19 | 11.52 | | | | | |
| Cortical depth | Deep | 1511.72 | 12.38 | 3 | 31 | 1148.357 | $4.760*10^{-43}$ | 0.743 |
| | Inner Middle | 1649.06 | 11.22 | | | | | |
| | Outer Middle | 1770.39 | 10.10 | | | | | |
| | Superficial | 1895.13 | 13.94 | | | | | |
| Distance | 2.01 - 4 mm | 1719.03 | 12.16 | 3 | 31 | 6.494 | 0.009 | 0.014 |
| | 4.01 - 6 mm | 1712.38 | 11.53 | | | | | |
| | 6.01 - 8 mm | 1701.47 | 10.89 | | | | | |
| | 8.01 - 10 mm | 1693.41 | 12.87 | | | | | |
| Age x Distance | Younger / 2.01 - 4 mm | 1740.50 | 16.95 | 3 | 31 | 19.232 | $1.447*10^{-5}$ | 0.040 |
| | Younger / 4.01 - 6 mm | 1728.96 | 16.06 | | | | | |
| | Younger / 6.01 - 8 mm | 1702.08 | 15.18 | | | | | |
| | Younger / 8.01 - 10 mm | 1671.42 | 17.94 | | | | | |
| | Older / 2.01 - 4 mm | 1697.57 | 17.44 | | | | | |
| | Older / 4.01 - 6 mm | 1695.80 | 16.53 | | | | | |
| | Older / 6.01 - 8 mm | 1700.87 | 15.62 | | | | | |
| | Older / 8.01 - 10 mm | 1715.39 | 18.46 | | | | | |
| Age x Brain Area | Younger / M1 | 1697.34 | 17.15 | 1 | 33 | 2.116 | 0.155 | 0.005 |
| | Older / M1 | 1682.57 | 17.64 | | | | | |
| | Younger / S1 | 1742.14 | 16.06 | | | | | |
| | Older / S1 | 1722.25 | 16.53 | | | | | |
| Brain Area x Distance | M1 / 2.01 - 4 mm | 1704.16 | 12.67 | 3 | 31 | 5.407 | 0.015 | 0.007 |
| | M1 / 4.01 - 6 mm | 1687.94 | 11.69 | | | | | |
| | M1 / 6.01 - 8 mm | 1671.56 | 11.81 | | | | | |
| | M1 / 8.01 - 10 mm | 1660.16 | 16.50 | | | | | |
| | S1 / 2.01 - 4 mm | 1733.91 | 12.96 | | | | | |
| | S1 / 4.01 - 6 mm | 1736.82 | 12.58 | | | | | |
| | S1 / 6.01 - 8 mm | 1731.39 | 11.47 | | | | | |
| | S1 / 8.01 - 10 mm | 1726.65 | 12.68 | | | | | |
| Age x Brain Area x Distance | Younger / M1 / 2.01 - 4 mm | 1725.63 | 17.66 | 3 | 31 | 3.714 | 0.024 | 0.005 |
| | Younger / M1 / 4.01 - 6 mm | 1702.89 | 16.30 | | | | | |
| | Younger / M1 / 6.01 - 8 mm | 1666.23 | 16.46 | | | | | |
| | Younger / M1 / 8.01 - 10 mm | 1622.62 | 23.00 | | | | | |
| | Younger / S1 / 2.01 - 4 mm | 1755.36 | 18.06 | | | | | |
| | Younger / S1 / 4.01 - 6 mm | 1755.03 | 17.53 | | | | | |
| | Younger / S1 / 6.01 - 8 mm | 1737.92 | 15.99 | | | | | |
| | Younger / S1 / 8.01 - 10 mm | 1720.23 | 17.68 | | | | | |
| | Older / M1 / 2.01 - 4 mm | 1682.68 | 18.18 | | | | | |
| | Older / M1 / 4.01 - 6 mm | 1672.99 | 16.77 | | | | | |
| | Older / M1 / 6.01 - 8 mm | 1676.88 | 16.94 | | | | | |
| | Older / M1 / 8.01 - 10 mm | 1697.71 | 23.66 | | | | | |
| | Older / S1 / 2.01 - 4 mm | 1712.45 | 18.58 | | | | | |
| | Older / S1 / 4.01 - 6 mm | 1718.60 | 18.04 | | | | | |
| | Older / S1 / 6.01 - 8 mm | 1724.86 | 16.45 | | | | | |
| | Older / S1 / 8.01 - 10 mm | 1733.07 | 18.19 | | | | | |

**Supplementary Table 3. ANOVA Results for qT1 values excluding the closest distance.** Results for the ANOVA on qT1 values with factors age (younger adults, older adults), cortical depth (SF, OM, IM, DP), brain area (S1, M1) and venous distance (4 distances, excluding the distance 0.01 - 2 mm). Given are mean qT1 values in milliseconds (Mean in ms), degrees of freedom of the numerator (DFn), degrees of freedom of the denominator (DFd), p-values (p), and effect sizes ($nG^2$). P-values below 0.05 are considered as significant.

| Effect | Conditions | Mean (ms) | SEM (ms) | Statistics | | | | |
|---|---|---|---|---|---|---|---|---|
| | | | | DFn | DFd | F | p | $\eta_G^2$ |
| Age | Younger | 1725.97 | 16.19 | 1 | 33 | 1.010 | 0.322 | 0.017 |
| | Older | 1702.26 | 17.17 | | | | | |
| Brain Area | M1 | 1692.69 | 13.11 | 1 | 33 | 23.004 | $3.590*10^{-5}$ | 0.055 |
| | S1 | 1735.54 | 12.11 | | | | | |
| Cortical depth | Deep | 1522.65 | 13.49 | 3 | 31 | 926.059 | $1.378*10^{-37}$ | 0.711 |
| | Inner Middle | 1657.41 | 12.00 | | | | | |
| | Outer Middle | 1776.73 | 10.53 | | | | | |
| | Superficial | 1899.67 | 14.30 | | | | | |
| Distance | 0.01 - 2 mm | 1745.46 | 13.79 | 4 | 30 | 25.229 | $3.039*10^{-8}$ | 0.044 |
| | 2.01 - 4 mm | 1721.05 | 12.38 | | | | | |
| | 4.01 - 6 mm | 1713.99 | 11.78 | | | | | |
| | 6.01 - 8 mm | 1700.61 | 11.20 | | | | | |
| | 8.01 - 10 mm | 1689.45 | 12.67 | | | | | |
| Age x Distance | Younger / 0.01 - 2 mm | 1786.91 | 18.92 | 4 | 30 | 26.928 | $1.286*10^{-8}$ | 0.047 |
| | Younger / 2.01 - 4 mm | 1740.50 | 16.99 | | | | | |
| | Younger / 4.01 - 6 mm | 1728.96 | 16.16 | | | | | |
| | Younger / 6.01 - 8 mm | 1702.08 | 15.36 | | | | | |
| | Younger / 8.01 - 10 mm | 1671.42 | 17.38 | | | | | |
| | Older / 0.01 - 2 mm | 1704.02 | 20.07 | | | | | |
| | Older / 2.01 - 4 mm | 1701.61 | 18.02 | | | | | |
| | Older / 4.01 - 6 mm | 1699.02 | 17.14 | | | | | |
| | Older / 6.01 - 8 mm | 1699.15 | 16.30 | | | | | |
| | Older / 8.01 - 10 mm | 1707.48 | 18.44 | | | | | |
| Age x Brain Area | Younger / M1 | 1700.53 | 17.98 | 1 | 33 | 0.808 | 0.376 | 0.002 |
| | Older / M1 | 1684.84 | 19.07 | | | | | |
| | Younger / S1 | 1751.41 | 16.61 | | | | | |
| | Older / S1 | 1719.67 | 17.62 | | | | | |
| Brain Area x Distance | M1 / 0.01 - 2 mm | 1739.60 | 16.64 | 4 | 30 | 7.734 | $4.802*10^{-4}$ | 0.011 |
| | M1 / 2.01 - 4 mm | 1705.64 | 12.99 | | | | | |
| | M1 / 4.01 - 6 mm | 1689.10 | 12.01 | | | | | |
| | M1 / 6.01 - 8 mm | 1670.83 | 12.16 | | | | | |
| | M1 / 8.01 - 10 mm | 1658.28 | 16.92 | | | | | |
| | S1 / 0.01 - 2 mm | 1751.33 | 14.83 | | | | | |
| | S1 / 2.01 - 4 mm | 1736.47 | 13.12 | | | | | |
| | S1 / 4.01 - 6 mm | 1738.88 | 12.81 | | | | | |
| | S1 / 6.01 - 8 mm | 1730.40 | 11.79 | | | | | |
| | S1 / 8.01 - 10 mm | 1720.63 | 11.61 | | | | | |
| Age x Brain Area x Distance | Younger / M1 / 0.01 - 2 mm | 1739.60 | 16.64 | 4 | 30 | 4.868 | 0.007 | 0.007 |
| | Younger / M1 / 2.01 - 4 mm | 1705.64 | 12.99 | | | | | |
| | Younger / M1 / 4.01 - 6 mm | 1689.10 | 12.01 | | | | | |
| | Younger / M1 / 6.01 - 8 mm | 1670.83 | 12.16 | | | | | |
| | Younger / M1 / 8.01 - 10 mm | 1658.28 | 16.92 | | | | | |
| | Younger / S1 / 0.01 - 2 mm | 1751.33 | 14.83 | | | | | |
| | Younger / S1 / 2.01 - 4 mm | 1736.47 | 13.12 | | | | | |
| | Younger / S1 / 4.01 - 6 mm | 1738.88 | 12.81 | | | | | |
| | Younger / S1 / 6.01 - 8 mm | 1730.40 | 11.79 | | | | | |
| | Younger / S1 / 8.01 - 10 mm | 1720.63 | 11.61 | | | | | |
| | Older / M1 / 0.01 - 2 mm | 1693.90 | 24.22 | | | | | |
| | Older / M1 / 2.01 - 4 mm | 1685.65 | 18.90 | | | | | |
| | Older / M1 / 4.01 - 6 mm | 1675.31 | 17.47 | | | | | |
| | Older / M1 / 6.01 - 8 mm | 1675.42 | 17.70 | | | | | |
| | Older / M1 / 8.01 - 10 mm | 1693.93 | 24.62 | | | | | |
| | Older / S1 / 0.01 - 2 mm | 1714.13 | 21.58 | | | | | |
| | Older / S1 / 2.01 - 4 mm | 1717.57 | 19.09 | | | | | |
| | Older / S1 / 4.01 - 6 mm | 1722.72 | 18.64 | | | | | |
| | Older / S1 / 6.01 - 8 mm | 1722.88 | 17.16 | | | | | |
| | Older / S1 / 8.01 - 10 mm | 1721.03 | 16.90 | | | | | |

**Supplementary Table 4. ANOVA Results for qT1 values without participant P2.** Results for the mixed-effects ANOVA on qT1 values with factors age (n=18 younger adults, n=17 older adults), cortical depth (SF, OM, IM, DP), brain area (S1, M1) and venous distance (5 distances). Given are mean qT1 values in milliseconds (Mean in ms) and standard error of the mean (SEM), degrees of freedom of the numerator (DFn), degrees of freedom of the denominator (DFd), p-values (p), and effect sizes ($n_G^2$). P-values ≤ 0.05 are considered as significant.

| condition | diff ± se | p | 95% CI |
|---|---|---|---|
| pQSM M1_0_2 | -0.0076 ± 0.0021 | $4*10^{-5}$ | -0.0118, -0.0034 |
| pQSM M1_2_4 | -0.0054 ± 0.0012 | $< 10^{-5}$ | -0.0078, -0.0030 |
| pQSM M1_4_6 | -0.0059 ± 0.0012 | $< 10^{-5}$ | -0.0083, -0.0036 |
| pQSM M1_6_8 | -0.0072 ± 0.0016 | $< 10^{-5}$ | -0.0103, -0.0041 |
| pQSM M1_8_10 | -0.0073 ± 0.0019 | $4*10^{-5}$ | -0.0110, -0.0036 |
| pQSM S1_0_2 | -0.0037 ± 0.0017 | 0.028 | -0.0069, -0.0004 |
| pQSM S1_2_4 | -0.0035 ± 0.0010 | $< 10^{-5}$ | -0.0053, -0.0016 |
| pQSM S1_4_6 | -0.0030 ± 0.0009 | 0.0001 | -0.0047, -0.0013 |
| pQSM S1_6_8 | -0.0031 ± 0.0008 | 0.034 | -0.0031, -0.0001 |
| pQSM S1_8_10 | -0.0011 ± 0.0011 | 0.295 | -0.0032, 0.0010 |

**Supplementary Table 5. Results of permutation Welch two sample t-tests comparing pQSM values between younger adults (n=18) and older adults (n=17).** Results are given as permutation mean difference ± standard error (diff ± se) in parts per million, permutation p-value (p) and 95% permutation confidence interval (95% CI).

| Effect | Conditions | Mean (ppm) | SEM (ppm) | Statistics | | | | |
|---|---|---|---|---|---|---|---|---|
| | | | | DFn | DFd | F | p | $\eta_G^2$ |
| Age | Younger | 0.0125 | 0.00047 | 1 | 33 | 44.722 | $1.507*10^{-7}$ | 0.261 |
| | Older | 0.0171 | 0.00050 | | | | | |
| Brain Area | M1 | 0.0163 | 0.00041 | 1 | 33 | 65.728 | $2.939*10^{-9}$ | 0.129 |
| | S1 | 0.0133 | 0.00037 | | | | | |
| Cortical depth | Deep | 0.0120 | 0.00033 | 3 | 31 | 143.172 | $1.598*10^{-23}$ | 0.166 |
| | Inner Middle | 0.0148 | 0.00037 | | | | | |
| | Outer Middle | 0.0163 | 0.00042 | | | | | |
| | Superficial | 0.0161 | 0.00037 | | | | | |
| Distance | 0.01 - 2 mm | 0.0162 | 0.00065 | 4 | 30 | 7.901 | 0.002 | 0.054 |
| | 2.01 - 4 mm | 0.0140 | 0.00038 | | | | | |
| | 4.01 - 6 mm | 0.0137 | 0.00030 | | | | | |
| | 6.01 - 8 mm | 0.0147 | 0.00036 | | | | | |
| | 8.01 - 10 mm | 0.0155 | 0.00059 | | | | | |
| Age x Distance | Younger / 0.01 - 2 mm | 0.0134 | 0.00089 | 4 | 30 | 0.629 | 0.500 | 0.005 |
| | Younger / 2.01 - 4 mm | 0.0118 | 0.00052 | | | | | |
| | Younger / 4.01 - 6 mm | 0.0115 | 0.00041 | | | | | |
| | Younger / 6.01 - 8 mm | 0.0125 | 0.00050 | | | | | |
| | Younger / 8.01 - 10 mm | 0.0134 | 0.00080 | | | | | |
| | Older / 0.01 - 2 mm | 0.0190 | 0.00094 | | | | | |
| | Older / 2.01 - 4 mm | 0.0162 | 0.00055 | | | | | |
| | Older / 4.01 - 6 mm | 0.0159 | 0.00043 | | | | | |
| | Older / 6.01 - 8 mm | 0.0168 | 0.00053 | | | | | |
| | Older / 8.01 - 10 mm | 0.0176 | 0.00085 | | | | | |
| Age x Brain Area | Younger / M1 | 0.0130 | 0.00056 | 1 | 33 | 31.846 | $3.056*10^{-6}$ | 0.067 |
| | Older / M1 | 0.0196 | 0.00059 | | | | | |
| | Younger / S1 | 0.0121 | 0.00051 | | | | | |
| | Older / S1 | 0.0146 | 0.00054 | | | | | |
| Brain Area x Distance | M1 / 0.01 - 2 mm | 0.0177 | 0.00089 | 4 | 30 | 0.224 | 0.756 | 0.001 |
| | M1 / 2.01 - 4 mm | 0.0157 | 0.00044 | | | | | |
| | M1 / 4.01 - 6 mm | 0.0152 | 0.00033 | | | | | |
| | M1 / 6.01 - 8 mm | 0.0162 | 0.00051 | | | | | |
| | M1 / 8.01 - 10 mm | 0.0167 | 0.00074 | | | | | |
| | S1 / 0.01 - 2 mm | 0.0147 | 0.00082 | | | | | |
| | S1 / 2.01 - 4 mm | 0.0124 | 0.00039 | | | | | |
| | S1 / 4.01 - 6 mm | 0.0122 | 0.00036 | | | | | |
| | S1 / 6.01 - 8 mm | 0.0131 | 0.00038 | | | | | |
| | S1 / 8.01 - 10 mm | 0.0143 | 0.00055 | | | | | |
| Age x Brain Area x Distance | Younger / M1 / 0.01 - 2 mm | 0.0138 | 0.00123 | 4 | 30 | 1.942 | 0.161 | 0.009 |
| | Younger / M1 / 2.01 - 4 mm | 0.0129 | 0.00060 | | | | | |
| | Younger / M1 / 4.01 - 6 mm | 0.0122 | 0.00046 | | | | | |
| | Younger / M1 / 6.01 - 8 mm | 0.0127 | 0.00070 | | | | | |
| | Younger / M1 / 8.01 - 10 mm | 0.0132 | 0.00102 | | | | | |
| | Younger / S1 / 0.01 - 2 mm | 0.0129 | 0.00113 | | | | | |
| | Younger / S1 / 2.01 - 4 mm | 0.0107 | 0.00053 | | | | | |
| | Younger / S1 / 4.01 - 6 mm | 0.0107 | 0.00050 | | | | | |
| | Younger / S1 / 6.01 - 8 mm | 0.0124 | 0.00052 | | | | | |
| | Younger / S1 / 8.01 - 10 mm | 0.0137 | 0.00076 | | | | | |
| | Older / M1 / 0.01 - 2 mm | 0.0215 | 0.00130 | | | | | |
| | Older / M1 / 2.01 - 4 mm | 0.0184 | 0.00063 | | | | | |
| | Older / M1 / 4.01 - 6 mm | 0.0181 | 0.00049 | | | | | |
| | Older / M1 / 6.01 - 8 mm | 0.0197 | 0.00074 | | | | | |
| | Older / M1 / 8.01 - 10 mm | 0.0203 | 0.00108 | | | | | |
| | Older / S1 / 0.01 - 2 mm | 0.0165 | 0.00120 | | | | | |
| | Older / S1 / 2.01 - 4 mm | 0.0141 | 0.00056 | | | | | |
| | Older / S1 / 4.01 - 6 mm | 0.0136 | 0.00053 | | | | | |
| | Older / S1 / 6.01 - 8 mm | 0.0139 | 0.00055 | | | | | |
| | Older / S1 / 8.01 - 10 mm | 0.0148 | 0.00080 | | | | | |

**Supplementary Table 6. ANOVA Results for pQSM values without participant P2.** Results for the ANOVA on pQSM values with factors age (n=18 younger adults, n=16 older adults), cortical depth (SF, OM, IM, DP), brain area (S1, M1) and venous distance (5 distances). Given are mean pQSM values in parts per million (Mean in ppm) and standard error of the mean (SEM), degrees of freedom of the numerator (DFn), degrees of freedom of the denominator (DFd), p-values (p), and effect sizes ($n_G^2$). P-values ≤ 0.05 are considered significant.

| Effect | Conditions | Mean (ppm) | SEM (ppm) | Statistics | | | | |
|---|---|---|---|---|---|---|---|---|
| | | | | DFn | DFd | F | p | $\eta_G^2$ |
| Age | Younger | 0.0123 | 0.00048 | 1 | 33 | 39.885 | $3.824*10^{-7}$ | 0.337 |
| | Older | 0.0167 | 0.00050 | | | | | |
| Brain Area | M1 | 0.0160 | 0.00042 | 1 | 33 | 78.835 | $2.912*10^{-10}$ | 0.189 |
| | S1 | 0.0130 | 0.00035 | | | | | |
| Cortical depth | Deep | 0.0115 | 0.00028 | 3 | 31 | 134.910 | $1.173*10^{-19}$ | 0.264 |
| | Inner Middle | 0.0144 | 0.00038 | | | | | |
| | Outer Middle | 0.0162 | 0.00045 | | | | | |
| | Superficial | 0.0158 | 0.00040 | | | | | |
| Distance | 2.01 - 4 mm | 0.0140 | 0.00037 | 3 | 31 | 10.687 | $4.384*10^{-4}$ | 0.050 |
| | 4.01 - 6 mm | 0.0137 | 0.00029 | | | | | |
| | 6.01 - 8 mm | 0.0147 | 0.00036 | | | | | |
| | 8.01 - 10 mm | 0.0155 | 0.00057 | | | | | |
| Age x Distance | Younger / 2.01 - 4 mm | 0.0118 | 0.00051 | 3 | 31 | 0.052 | 0.911 | $2.525*10^{-4}$ |
| | Younger / 4.01 - 6 mm | 0.0115 | 0.00040 | | | | | |
| | Younger / 6.01 - 8 mm | 0.0125 | 0.00050 | | | | | |
| | Younger / 8.01 - 10 mm | 0.0134 | 0.00079 | | | | | |
| | Older / 2.01 - 4 mm | 0.0163 | 0.00052 | | | | | |
| | Older / 4.01 - 6 mm | 0.0159 | 0.00042 | | | | | |
| | Older / 6.01 - 8 mm | 0.0169 | 0.00051 | | | | | |
| | Older / 8.01 - 10 mm | 0.0177 | 0.00082 | | | | | |
| Age x Brain Area | Younger / M1 | 0.0128 | 0.00058 | 1 | 33 | 38.484 | $5.306*10^{-7}$ | 0.102 |
| | Older / M1 | 0.0192 | 0.00060 | | | | | |
| | Younger / S1 | 0.0119 | 0.00049 | | | | | |
| | Older / S1 | 0.0142 | 0.00051 | | | | | |
| Brain Area x Distance | M1 / 2.01 - 4 mm | 0.0156 | 0.00042 | 3 | 31 | 0.695 | 0.485 | 0.002 |
| | M1 / 4.01 - 6 mm | 0.0152 | 0.00032 | | | | | |
| | M1 / 6.01 - 8 mm | 0.0163 | 0.00050 | | | | | |
| | M1 / 8.01 - 10 mm | 0.0168 | 0.00072 | | | | | |
| | S1 / 2.01 - 4 mm | 0.0124 | 0.00038 | | | | | |
| | S1 / 4.01 - 6 mm | 0.0122 | 0.00035 | | | | | |
| | S1 / 6.01 - 8 mm | 0.0132 | 0.00037 | | | | | |
| | S1 / 8.01 - 10 mm | 0.0143 | 0.00053 | | | | | |
| Age x Brain Area x Distance | Younger / M1 / 2.01 - 4 mm | 0.0129 | 0.00059 | 3 | 31 | 8.767 | 0.001 | 0.020 |
| | Younger / M1 / 4.01 - 6 mm | 0.0122 | 0.00045 | | | | | |
| | Younger / M1 / 6.01 - 8 mm | 0.0127 | 0.00070 | | | | | |
| | Younger / M1 / 8.01 - 10 mm | 0.0132 | 0.00101 | | | | | |
| | Younger / S1 / 2.01 - 4 mm | 0.0107 | 0.00052 | | | | | |
| | Younger / S1 / 4.01 - 6 mm | 0.0107 | 0.00049 | | | | | |
| | Younger / S1 / 6.01 - 8 mm | 0.0124 | 0.00051 | | | | | |
| | Younger / S1 / 8.01 - 10 mm | 0.0137 | 0.00074 | | | | | |
| | Older / M1 / 2.01 - 4 mm | 0.0183 | 0.00061 | | | | | |
| | Older / M1 / 4.01 - 6 mm | 0.0182 | 0.00046 | | | | | |
| | Older / M1 / 6.01 - 8 mm | 0.0199 | 0.00072 | | | | | |
| | Older / M1 / 8.01 - 10 mm | 0.0205 | 0.00104 | | | | | |
| | Older / S1 / 2.01 - 4 mm | 0.0142 | 0.00054 | | | | | |
| | Older / S1 / 4.01 - 6 mm | 0.0137 | 0.00051 | | | | | |
| | Older / S1 / 6.01 - 8 mm | 0.0140 | 0.00053 | | | | | |
| | Older / S1 / 8.01 - 10 mm | 0.0148 | 0.00077 | | | | | |

**Supplementary Table 7. ANOVA Results for pQSM values excluding the closest distance.** Results for the ANOVA on pQSM values with factors age (younger adults, older adults), cortical depth (SF, OM, IM, DP), brain area (S1, M1) and venous distance (4 distances, excluding the distance 0.01 - 2 mm). Given are mean pQSM values in parts per million (Mean in ppm), degrees of freedom of the numerator (DFn), degrees of freedom of the denominator (DFd), p-values (p), and effect sizes ($nG^2$). P-values below 0.05 are considered as significant.

| condition | younger adults | older adults |
| --- | --- | --- |
| M1 0-2 mm | -0.24 | -0.21 |
| M1 2-4 mm | **-0.46** | -0.21 |
| M1 4-6 mm | **-0.38** | 0.00 |
| M1 6-8 mm | 0.05 | -0.04 |
| M1 8-10 mm | 0.11 | -0.14 |
| S1 0-2 mm | -0.13 | -0.24 |
| S1 2-4 mm | **-0.31** | **-0.41** |
| S1 4-6 mm | -0.28 | -0.19 |
| S1 6-8 mm | **-0.37** | -0.18 |
| S1 8-10 mm | -0.21 | -0.19 |

**Supplementary Table 8. Pearson correlation coefficients given for the correlation between qT1 and pQSM values for younger adults (n=18) and older adults (n=17) for each area-by-distance condition (as shown in Supplementary Figure 3).** Areas: primary motor cortex (M1), primary somatosensory cortex (S1). Venous distances in millimetres (mm): 0-2 mm, 2-4 mm, 4-6 mm, 6-8 mm, 8-10 mm. Small correlation: $r < 0.3$, moderate correlation: $r \geq 0.3$ (highlighted in bold), strong correlation: $r \geq 0.5$.

| condition | diff ± se | p | 95% CI |
|---|---|---|---|
| nQSM M1_0_2 | 0.0040 ± 0.0014 | 0.003 | 0.0012, 0.0068 |
| nQSM M1_2_4 | 0.0030 ± 0.0008 | $< 10^{-5}$ | 0.0015, 0.0044 |
| nQSM M1_4_6 | 0.0024 ± 0.0006 | $< 10^{-5}$ | 0.0012, 0.0036 |
| nQSM M1_6_8 | 0.0025 ± 0.0007 | 0.0001 | 0.0011, 0.0039 |
| nQSM M1_8_10 | 0.0019 ± 0.0010 | 0.062 | -0.0001, 0.0039 |
| nQSM S1_0_2 | 0.0032 ± 0.0012 | 0.005 | 0.0009, 0.0056 |
| nQSM S1_2_4 | 0.0022 ± 0.0007 | 0.001 | 0.0008, 0.0036 |
| nQSM S1_4_6 | 0.0021 ± 0.0006 | 0.0002 | 0.0009, 0.0032 |
| nQSM S1_6_8 | 0.0029 ± 0.0007 | $4*10^{-5}$ | 0.0015, 0.0043 |
| nQSM S1_8_10 | 0.0031 ± 0.0009 | 0.0001 | 0.0014, 0.0049 |

**Supplementary Table 9. Results of permutation Welch two sample t-tests comparing nQSM values between younger adults (n=18) and older adults (n=17).** Results are given as permutation mean difference ± standard error (diff ± se) in parts per million, permutation p-value (p) and 95% permutation confidence interval (95% CI).

| Effect | Conditions | Mean (ppm) | SEM (ppm) | Statistics DFn | DFd | F | p | $\eta_G^2$ |
|---|---|---|---|---|---|---|---|---|
| Age | Younger | -0.0095 | 0.00038 | 1 | 33 | 25.136 | $1.772*10^{-5}$ | 0.177 |
| | Older | -0.0122 | 0.00039 | | | | | |
| Brain Area | M1 | -0.0105 | 0.00031 | 1 | 33 | 11.737 | 0.002 | 0.016 |
| | S1 | -0.0113 | 0.00027 | | | | | |
| Cortical depth | Deep | -0.0123 | 0.00033 | 3 | 31 | 49.416 | $1.068*10^{-10}$ | 0.162 |
| | Inner Middle | -0.0097 | 0.00033 | | | | | |
| | Outer Middle | -0.0095 | 0.00031 | | | | | |
| | Superficial | -0.0121 | 0.00033 | | | | | |
| Distance | 0.01 - 2 mm | -0.0115 | 0.00054 | 4 | 30 | 2.048 | 0.138 | 0.011 |
| | 2.01 - 4 mm | -0.0108 | 0.00027 | | | | | |
| | 4.01 - 6 mm | -0.0108 | 0.00019 | | | | | |
| | 6.01 - 8 mm | -0.0108 | 0.00025 | | | | | |
| | 8.01 - 10 mm | -0.0106 | 0.00037 | | | | | |
| Age x Distance | Younger / 0.01 - 2 mm | -0.0097 | 0.00075 | 4 | 30 | 1.215 | 0.303 | 0.007 |
| | Younger / 2.01 - 4 mm | -0.0095 | 0.00037 | | | | | |
| | Younger / 4.01 - 6 mm | -0.0097 | 0.00026 | | | | | |
| | Younger / 6.01 - 8 mm | -0.0095 | 0.00035 | | | | | |
| | Younger / 8.01 - 10 mm | -0.0093 | 0.00051 | | | | | |
| | Older / 0.01 - 2 mm | -0.0133 | 0.00078 | | | | | |
| | Older / 2.01 - 4 mm | -0.0120 | 0.00038 | | | | | |
| | Older / 4.01 - 6 mm | -0.0119 | 0.00027 | | | | | |
| | Older / 6.01 - 8 mm | -0.0121 | 0.00036 | | | | | |
| | Older / 8.01 - 10 mm | -0.0118 | 0.00053 | | | | | |
| Age x Brain Area | Younger / M1 | -0.0091 | 0.00043 | 1 | 33 | 0.008 | 0.928 | $1*10^{-5}$ |
| | Older / M1 | -0.0099 | 0.00038 | | | | | |
| | Younger / S1 | -0.0119 | 0.00045 | | | | | |
| | Older / S1 | -0.0126 | 0.00039 | | | | | |
| Brain Area x Distance | M1 / 0.01 - 2 mm | -0.0101 | 0.00065 | 4 | 30 | 13.493 | $1.296*10^{-6}$ | 0.037 |
| | M1 / 2.01 - 4 mm | -0.0102 | 0.00028 | | | | | |
| | M1 / 4.01 - 6 mm | -0.0105 | 0.00023 | | | | | |
| | M1 / 6.01 - 8 mm | -0.0108 | 0.00030 | | | | | |
| | M1 / 8.01 - 10 mm | -0.0109 | 0.00049 | | | | | |
| | S1 / 0.01 - 2 mm | -0.0129 | 0.00056 | | | | | |
| | S1 / 2.01 - 4 mm | -0.0113 | 0.00032 | | | | | |
| | S1 / 4.01 - 6 mm | -0.0111 | 0.00024 | | | | | |
| | S1 / 6.01 - 8 mm | -0.0108 | 0.00027 | | | | | |
| | S1 / 8.01 - 10 mm | -0.0103 | 0.00037 | | | | | |
| Age x Brain Area x Distance | Younger / M1 / 0.01 - 2 mm | -0.0081 | 0.00091 | 4 | 30 | 1.506 | 0.224 | 0.004 |
| | Younger / M1 / 2.01 - 4 mm | -0.0088 | 0.00039 | | | | | |
| | Younger / M1 / 4.01 - 6 mm | -0.0093 | 0.00032 | | | | | |
| | Younger / M1 / 6.01 - 8 mm | -0.0096 | 0.00042 | | | | | |
| | Younger / M1 / 8.01 - 10 mm | -0.0100 | 0.00068 | | | | | |
| | Younger / S1 / 0.01 - 2 mm | -0.0112 | 0.00077 | | | | | |
| | Younger / S1 / 2.01 - 4 mm | -0.0102 | 0.00045 | | | | | |
| | Younger / S1 / 4.01 - 6 mm | -0.0100 | 0.00033 | | | | | |
| | Younger / S1 / 6.01 - 8 mm | -0.0094 | 0.00038 | | | | | |
| | Younger / S1 / 8.01 - 10 mm | -0.0087 | 0.00051 | | | | | |
| | Older / M1 / 0.01 - 2 mm | -0.0121 | 0.00093 | | | | | |
| | Older / M1 / 2.01 - 4 mm | -0.0117 | 0.00040 | | | | | |
| | Older / M1 / 4.01 - 6 mm | -0.0117 | 0.00033 | | | | | |
| | Older / M1 / 6.01 - 8 mm | -0.0120 | 0.00043 | | | | | |
| | Older / M1 / 8.01 - 10 mm | -0.0118 | 0.00070 | | | | | |
| | Older / S1 / 0.01 - 2 mm | -0.0145 | 0.00080 | | | | | |
| | Older / S1 / 2.01 - 4 mm | -0.0124 | 0.00046 | | | | | |
| | Older / S1 / 4.01 - 6 mm | -0.0121 | 0.00034 | | | | | |
| | Older / S1 / 6.01 - 8 mm | -0.0123 | 0.00039 | | | | | |
| | Older / S1 / 8.01 - 10 mm | -0.0118 | 0.00052 | | | | | |

**Supplementary Table 10. ANOVA Results for nQSM values.** Results for the ANOVA on nQSM values with factors age (n=18 younger adults, n=17 older adults), cortical depth (SF, OM, IM, DP), brain area (S1, M1) and venous distance (5 distances). Given are mean nQSM values in parts per million (Mean in ppm) and standard error of the mean (SEM), degrees of freedom of the numerator (DFn), degrees of freedom of the denominator (DFd), p-values (p), and effect sizes ($n_G^2$). P-values $\leq$ 0.05 are considered as significant.

| Effect | Conditions | Mean | SEM | Statistics | | | | |
|---|---|---|---|---|---|---|---|---|
| | | (ppm) | (ppm) | DFn | DFd | F | p | $\eta_G^2$ |
| Age | Younger | -0.00949 | 0.00033 | 1 | 33 | 28.243 | $7.295*10^{-6}$ | 0.190 |
| | Older | -0.01198 | 0.00034 | | | | | |
| Brain Area | M1 | -0.01061 | 0.00028 | 1 | 33 | 1.199 | 0.282 | 0.002 |
| | S1 | -0.01086 | 0.00024 | | | | | |
| Cortical depth | Deep | -0.01296 | 0.00033 | 3 | 31 | 60.157 | $9.068*10^{-12}$ | 0.254 |
| | Inner Middle | -0.00981 | 0.00031 | | | | | |
| | Outer Middle | -0.00900 | 0.00027 | | | | | |
| | Superficial | -0.01116 | 0.00031 | | | | | |
| Distance | 2.01 - 4 mm | -0.01075 | 0.00027 | 3 | 31 | 0.389 | 0.656 | 0.001 |
| | 4.01 - 6 mm | -0.01079 | 0.00019 | | | | | |
| | 6.01 - 8 mm | -0.01082 | 0.00025 | | | | | |
| | 8.01 - 10 mm | -0.01058 | 0.00037 | | | | | |
| Age x Distance | Younger / 2.01 - 4 mm | -0.00946 | 0.00037 | 3 | 31 | 0.328 | 0.697 | 0.001 |
| | Younger / 4.01 - 6 mm | -0.00968 | 0.00026 | | | | | |
| | Younger / 6.01 - 8 mm | -0.00948 | 0.00035 | | | | | |
| | Younger / 8.01 - 10 mm | -0.00933 | 0.00051 | | | | | |
| | Older / 2.01 - 4 mm | -0.01204 | 0.00038 | | | | | |
| | Older / 4.01 - 6 mm | -0.01190 | 0.00027 | | | | | |
| | Older / 6.01 - 8 mm | -0.01215 | 0.00036 | | | | | |
| | Older / 8.01 - 10 mm | -0.01183 | 0.00053 | | | | | |
| Age x Brain Area | Younger / M1 | -0.00940 | 0.00039 | 1 | 33 | 0.105 | 0.748 | $2*10^{-4}$ |
| | Older / M1 | -0.00958 | 0.00033 | | | | | |
| | Younger / S1 | -0.01182 | 0.00040 | | | | | |
| | Older / S1 | -0.01214 | 0.00034 | | | | | |
| Brain Area x Distance | M1 / 2.01 - 4 mm | -0.01024 | 0.00028 | 3 | 31 | 6.589 | 0.003 | 0.014 |
| | M1 / 4.01 - 6 mm | -0.01050 | 0.00023 | | | | | |
| | M1 / 6.01 - 8 mm | -0.01080 | 0.00030 | | | | | |
| | M1 / 8.01 - 10 mm | -0.01090 | 0.00049 | | | | | |
| | S1 / 2.01 - 4 mm | -0.01126 | 0.00032 | | | | | |
| | S1 / 4.01 - 6 mm | -0.01108 | 0.00024 | | | | | |
| | S1 / 6.01 - 8 mm | -0.01083 | 0.00027 | | | | | |
| | S1 / 8.01 - 10 mm | -0.01026 | 0.00037 | | | | | |
| Age x Brain Area x Distance | Younger / M1 / 2.01 - 4 mm | -0.00876 | 0.00039 | 3 | 31 | 2.410 | 0.098 | 0.005 |
| | Younger / M1 / 4.01 - 6 mm | -0.00931 | 0.00032 | | | | | |
| | Younger / M1 / 6.01 - 8 mm | -0.00958 | 0.00042 | | | | | |
| | Younger / M1 / 8.01 - 10 mm | -0.00996 | 0.00068 | | | | | |
| | Younger / S1 / 2.01 - 4 mm | -0.01016 | 0.00045 | | | | | |
| | Younger / S1 / 4.01 - 6 mm | -0.01005 | 0.00033 | | | | | |
| | Younger / S1 / 6.01 - 8 mm | -0.00939 | 0.00038 | | | | | |
| | Younger / S1 / 8.01 - 10 mm | -0.00871 | 0.00051 | | | | | |
| | Older / M1 / 2.01 - 4 mm | -0.01172 | 0.00040 | | | | | |
| | Older / M1 / 4.01 - 6 mm | -0.01168 | 0.00033 | | | | | |
| | Older / M1 / 6.01 - 8 mm | -0.01203 | 0.00043 | | | | | |
| | Older / M1 / 8.01 - 10 mm | -0.01185 | 0.00070 | | | | | |
| | Older / S1 / 2.01 - 4 mm | -0.01236 | 0.00046 | | | | | |
| | Older / S1 / 4.01 - 6 mm | -0.01212 | 0.00034 | | | | | |
| | Older / S1 / 6.01 - 8 mm | -0.01227 | 0.00039 | | | | | |
| | Older / S1 / 8.01 - 10 mm | -0.01182 | 0.00052 | | | | | |

**Supplementary Table 11. ANOVA Results for nQSM values excluding the closest distance.** Results for the ANOVA on nQSM values with factors age (younger adults, older adults), cortical depth (SF, OM, IM, DP), brain area (S1, M1) and venous distance (4 distances, excluding the distance 0.01 - 2 mm). Given are mean nQSM values in parts per million (Mean in ppm), degrees of freedom of the numerator (DFn), degrees of freedom of the denominator (DFd), p-values (p), and effect sizes ($nG^2$). P-values below 0.05 are considered as significant.

| | Conditions | Mean | SEM | Statistics | | | | |
|---|---|---|---|---|---|---|---|---|
| | | (ppm) | (ppm) | DFn | DFd | F | p | $\eta_G^2$ |
| Age | Younger | 1819.41 | 14.70 | 1 | 33 | 1.389 | 0.247 | 0.024 |
| | Older | 1794.55 | 15.13 | | | | | |
| Brain Area | M1 | 1692.42 | 12.71 | 4 | 132 | 289.016 | $2.274*10^{-37}$ | 0.490 |
| | S1 | 1735.63 | 11.73 | | | | | |
| | SFG | 1884.02 | 10.43 | | | | | |
| | CMF | 1877.92 | 10.80 | | | | | |
| | RMF | 1844.91 | 11.73 | | | | | |
| Cortical depth | Deep | 1600.74 | 11.69 | 3 | 99 | 1961.016 | $3.481*10^{-50}$ | 0.750 |
| | Inner Middle | 1771.46 | 11.06 | | | | | |
| | Outer Middle | 1884.28 | 9.64 | | | | | |
| | Superficial | 1971.44 | 11.48 | | | | | |
| Distance | 0.01 - 2 mm | 1829.95 | 11.52 | 4 | 132 | 42.205 | $8.142*10^{-12}$ | 0.034 |
| | 2.01 - 4 mm | 1813.58 | 10.62 | | | | | |
| | 4.01 - 6 mm | 1808.50 | 10.37 | | | | | |
| | 6.01 - 8 mm | 1796.77 | 10.41 | | | | | |
| | 8.01 - 10 mm | 1786.11 | 11.02 | | | | | |
| Age x Distance | Younger / 0.01 - 2 mm | 1857.45 | 16.05 | 4 | 132 | 24.494 | $3.155*10^{-8}$ | 0.020 |
| | Younger / 2.01 - 4 mm | 1832.37 | 14.81 | | | | | |
| | Younger / 4.01 - 6 mm | 1823.99 | 14.46 | | | | | |
| | Younger / 6.01 - 8 mm | 1802.86 | 14.50 | | | | | |
| | Younger / 8.01 - 10 mm | 1780.40 | 15.36 | | | | | |
| | Older / 0.01 - 2 mm | 1802.45 | 16.52 | | | | | |
| | Older / 2.01 - 4 mm | 1794.80 | 15.24 | | | | | |
| | Older / 4.01 - 6 mm | 1793.02 | 14.88 | | | | | |
| | Older / 6.01 - 8 mm | 1790.67 | 14.92 | | | | | |
| | Older / 8.01 - 10 mm | 1791.82 | 15.81 | | | | | |
| Age x Brain Area | Younger / M1 | 1700.53 | 17.71 | 4 | 132 | 1.578 | 0.184 | 0.005 |
| | Older / M1 | 1751.41 | 16.36 | | | | | |
| | Younger / S1 | 1898.83 | 14.54 | | | | | |
| | Older / S1 | 1897.74 | 15.06 | | | | | |
| | Younger / SFG | 1848.56 | 16.35 | | | | | |
| | Older / SFG | 1684.31 | 18.22 | | | | | |
| | Younger / CMF | 1719.85 | 16.83 | | | | | |
| | Older / CMF | 1869.22 | 14.96 | | | | | |
| | Younger / RMF | 1858.12 | 15.50 | | | | | |
| | Older / RMF | 1841.28 | 16.83 | | | | | |
| Brain Area x Distance | M1 / 0.01 - 2 mm | 1738.28 | 16.18 | 16 | 528 | 4.045 | $9.324*10^{-4}$ | 0.010 |
| | M1 / 2.01 - 4 mm | 1704.16 | 12.67 | | | | | |
| | M1 / 4.01 - 6 mm | 1687.94 | 11.69 | | | | | |
| | M1 / 6.01 - 8 mm | 1671.56 | 11.81 | | | | | |
| | M1 / 8.01 - 10 mm | 1660.16 | 16.50 | | | | | |
| | S1 / 0.01 - 2 mm | 1749.39 | 14.49 | | | | | |
| | S1 / 2.01 - 4 mm | 1733.91 | 12.96 | | | | | |
| | S1 / 4.01 - 6 mm | 1736.82 | 12.58 | | | | | |
| | S1 / 6.01 - 8 mm | 1731.39 | 11.47 | | | | | |
| | S1 / 8.01 - 10 mm | 1726.65 | 12.68 | | | | | |
| | SFG / 0.01 - 2 mm | 1903.67 | 12.30 | | | | | |
| | SFG / 2.01 - 4 mm | 1892.39 | 10.85 | | | | | |
| | SFG / 4.01 - 6 mm | 1886.74 | 10.09 | | | | | |
| | SFG / 6.01 - 8 mm | 1873.53 | 10.75 | | | | | |
| | SFG / 8.01 - 10 mm | 1863.79 | 12.08 | | | | | |
| | CMF / 0.01 - 2 mm | 1892.05 | 12.50 | | | | | |
| | CMF / 2.01 - 4 mm | 1890.35 | 10.20 | | | | | |
| | CMF / 4.01 - 6 mm | 1887.24 | 10.53 | | | | | |
| | CMF / 6.01 - 8 mm | 1871.89 | 11.52 | | | | | |
| | CMF / 8.01 - 10 mm | 1848.10 | 11.93 | | | | | |
| | RMF / 0.01 - 2 mm | 1866.37 | 13.74 | | | | | |
| | RMF / 2.01 - 4 mm | 1847.11 | 11.10 | | | | | |
| | RMF / 4.01 - 6 mm | 1843.78 | 11.21 | | | | | |
| | RMF / 6.01 - 8 mm | 1835.47 | 11.76 | | | | | |
| | RMF / 8.01 - 10 mm | 1831.86 | 13.04 | | | | | |
| Age x Brain Area x Distance | Younger / M1 / 0.01 - 2 mm | 1785.29 | 22.55 | 16 | 528 | 7.198 | $9.514*10^{-7}$ | 0.017 |
| | Younger / M1 / 2.01 - 4 mm | 1725.63 | 17.66 | | | | | |
| | Younger / M1 / 4.01 - 6 mm | 1702.89 | 16.30 | | | | | |
| | Younger / M1 / 6.01 - 8 mm | 1666.23 | 16.46 | | | | | |
| | Younger / M1 / 8.01 - 10 mm | 1622.62 | 23.00 | | | | | |
| | Younger / S1 / 0.01 - 2 mm | 1788.53 | 20.20 | | | | | |
| | Younger / S1 / 2.01 - 4 mm | 1755.36 | 18.06 | | | | | |
| | Younger / S1 / 4.01 - 6 mm | 1755.03 | 17.53 | | | | | |
| | Younger / S1 / 6.01 - 8 mm | 1737.92 | 15.99 | | | | | |
| | Younger / S1 / 8.01 - 10 mm | 1720.23 | 17.68 | | | | | |
| | Younger / SFG / 0.01 - 2 mm | 1910.30 | 17.15 | | | | | |
| | Younger / SFG / 2.01 - 4 mm | 1911.40 | 15.13 | | | | | |
| | Younger / SFG / 4.01 - 6 mm | 1904.02 | 14.07 | | | | | |
| | Younger / SFG / 6.01 - 8 mm | 1889.34 | 14.98 | | | | | |
| | Younger / SFG / 8.01 - 10 mm | 1879.09 | 16.84 | | | | | |
| | Younger / CMF / 0.01 - 2 mm | 1924.67 | 17.42 | | | | | |
| | Younger / CMF / 2.01 - 4 mm | 1917.75 | 14.21 | | | | | |
| | Younger / CMF / 4.01 - 6 mm | 1911.02 | 14.68 | | | | | |

| | | |
|---|---|---|
| Younger / CMF / 6.01 - 8 mm | 1884.98 | 16.06 |
| Younger / CMF / 8.01 - 10 mm | 1850.27 | 16.63 |
| Younger / RMF / 0.01 - 2 mm | 1878.46 | 19.16 |
| Younger / RMF / 2.01 - 4 mm | 1851.70 | 15.48 |
| Younger / RMF / 4.01 - 6 mm | 1847.01 | 15.63 |
| Younger / RMF / 6.01 - 8 mm | 1835.84 | 16.39 |
| Younger / RMF / 8.01 - 10 mm | 1829.80 | 18.18 |
| Older / M1 / 0.01 - 2 mm | 1691.27 | 23.21 |
| Older / M1 / 2.01 - 4 mm | 1682.68 | 18.17 |
| Older / M1 / 4.01 - 6 mm | 1672.99 | 16.77 |
| Older / M1 / 6.01 - 8 mm | 1676.88 | 16.94 |
| Older / M1 / 8.01 - 10 mm | 1697.71 | 23.66 |
| Older / S1 / 0.01 - 2 mm | 1710.25 | 20.79 |
| Older / S1 / 2.01 - 4 mm | 1712.45 | 18.58 |
| Older / S1 / 4.01 - 6 mm | 1718.60 | 18.04 |
| Older / S1 / 6.01 - 8 mm | 1724.86 | 16.45 |
| Older / S1 / 8.01 - 10 mm | 1733.07 | 18.19 |
| Older / SFG / 0.01 - 2 mm | 1897.04 | 17.65 |
| Older / SFG / 2.01 - 4 mm | 1873.38 | 15.56 |
| Older / SFG / 4.01 - 6 mm | 1869.45 | 14.48 |
| Older / SFG / 6.01 - 8 mm | 1857.72 | 15.41 |
| Older / SFG / 8.01 - 10 mm | 1848.49 | 17.33 |
| Older / CMF / 0.01 - 2 mm | 1859.42 | 17.92 |
| Older / CMF / 2.01 - 4 mm | 1862.96 | 14.62 |
| Older / CMF / 4.01 - 6 mm | 1863.47 | 15.11 |
| Older / CMF / 6.01 - 8 mm | 1858.80 | 16.53 |
| Older / CMF / 8.01 - 10 mm | 1845.93 | 17.11 |
| Older / RMF / 0.01 - 2 mm | 1854.28 | 19.71 |
| Older / RMF / 2.01 - 4 mm | 1842.52 | 15.92 |
| Older / RMF / 4.01 - 6 mm | 1840.55 | 16.08 |
| Older / RMF / 6.01 - 8 mm | 1835.10 | 16.86 |
| Older / RMF / 8.01 - 10 mm | 1833.92 | 18.70 |

**Supplementary Table 12. ANOVA Results for qT1 values including the primary motor cortex (M1), primary somatosensory cortex (S1), superior frontal gyrus (SFG), caudal middle frontal cortex (CMF), and rostral middle frontal cortex (RMF).** Results for the ANOVA on qT1 values with factors age (n=18 younger adults, n=17 older adults), cortical depth (SF, OM, IM, DP), brain area (S1, M1, SFG, CMF, RMF) and venous distance (5 distances). Given are mean qT1 values in milliseconds (Mean in ms) and standard error of the mean (SEM), degrees of freedom of the numerator (DFn), degrees of freedom of the denominator (DFd), p-values (p), and effect sizes ($n_G^2$). P-values $\leq 0.05$ are considered as significant.

| | Conditions | Mean | SEM | Statistics | | | | |
|---|---|---|---|---|---|---|---|---|
| | | (ppm) | (ppm) | DFn | DFd | F | p | $\eta_G^2$ |
| Age | Younger | 0.01005 | 0.00036 | 1 | 33 | 51.273 | $3.321*10^{-8}$ | 0.246 |
| | Older | 0.01370 | 0.00037 | | | | | |
| Brain Area | M1 | 0.01631 | 0.00039 | 4 | 132 | 130.393 | $1.200*10^{-32}$ | 0.402 |
| | S1 | 0.01336 | 0.00036 | | | | | |
| | SFG | 0.00931 | 0.00026 | | | | | |
| | CMF | 0.00992 | 0.00039 | | | | | |
| | RMF | 0.01048 | 0.00028 | | | | | |
| Cortical depth | Deep | 0.01090 | 0.00027 | 3 | 99 | 28.519 | $2.896*10^{-8}$ | 0.035 |
| | Inner Middle | 0.01186 | 0.00027 | | | | | |
| | Outer Middle | 0.01225 | 0.00029 | | | | | |
| | Superficial | 0.01250 | 0.00028 | | | | | |
| Distance | 0.01 - 2 mm | 0.01317 | 0.00045 | 4 | 132 | 16.256 | $2.231*10^{-5}$ | 0.047 |
| | 2.01 - 4 mm | 0.01145 | 0.00027 | | | | | |
| | 4.01 - 6 mm | 0.01113 | 0.00022 | | | | | |
| | 6.01 - 8 mm | 0.01155 | 0.00023 | | | | | |
| | 8.01 - 10 mm | 0.01208 | 0.00032 | | | | | |
| Age x Distance | Younger / 0.01 - 2 mm | 0.01058 | 0.00062 | 4 | 132 | 5.170 | 0.015 | 0.016 |
| | Younger / 2.01 - 4 mm | 0.00964 | 0.00038 | | | | | |
| | Younger / 4.01 - 6 mm | 0.00944 | 0.00031 | | | | | |
| | Younger / 6.01 - 8 mm | 0.01000 | 0.00032 | | | | | |
| | Younger / 8.01 - 10 mm | 0.01062 | 0.00045 | | | | | |
| | Older / 0.01 - 2 mm | 0.01575 | 0.00064 | | | | | |
| | Older / 2.01 - 4 mm | 0.01326 | 0.00039 | | | | | |
| | Older / 4.01 - 6 mm | 0.01283 | 0.00032 | | | | | |
| | Older / 6.01 - 8 mm | 0.01310 | 0.00033 | | | | | |
| | Older / 8.01 - 10 mm | 0.01354 | 0.00046 | | | | | |
| Age x Brain Area | Younger / M1 | 0.01297 | 0.00055 | 4 | 132 | 11.941 | $1.689*10^{-6}$ | 0.058 |
| | Older / M1 | 0.01207 | 0.00050 | | | | | |
| | Younger / S1 | 0.00772 | 0.00037 | | | | | |
| | Older / S1 | 0.00816 | 0.00054 | | | | | |
| | Younger / SFG | 0.00935 | 0.00039 | | | | | |
| | Older / SFG | 0.01966 | 0.00056 | | | | | |
| | Younger / CMF | 0.01465 | 0.00052 | | | | | |
| | Older / CMF | 0.01091 | 0.00038 | | | | | |
| | Younger / RMF | 0.01168 | 0.00056 | | | | | |
| | Older / RMF | 0.01160 | 0.00040 | | | | | |
| Brain Area x Distance | M1 / 0.01 - 2 mm | 0.01763 | 0.00087 | 16 | 528 | 1.944 | 0.093 | 0.014 |
| | M1 / 2.01 - 4 mm | 0.01564 | 0.00042 | | | | | |
| | M1 / 4.01 - 6 mm | 0.01521 | 0.00032 | | | | | |
| | M1 / 6.01 - 8 mm | 0.01627 | 0.00050 | | | | | |
| | M1 / 8.01 - 10 mm | 0.01682 | 0.00072 | | | | | |
| | S1 / 0.01 - 2 mm | 0.01474 | 0.00080 | | | | | |
| | S1 / 2.01 - 4 mm | 0.01244 | 0.00038 | | | | | |
| | S1 / 4.01 - 6 mm | 0.01218 | 0.00035 | | | | | |
| | S1 / 6.01 - 8 mm | 0.01316 | 0.00037 | | | | | |
| | S1 / 8.01 - 10 mm | 0.01426 | 0.00053 | | | | | |
| | SFG / 0.01 - 2 mm | 0.01073 | 0.00048 | | | | | |
| | SFG / 2.01 - 4 mm | 0.00947 | 0.00029 | | | | | |
| | SFG / 4.01 - 6 mm | 0.00923 | 0.00024 | | | | | |
| | SFG / 6.01 - 8 mm | 0.00865 | 0.00023 | | | | | |
| | SFG / 8.01 - 10 mm | 0.00847 | 0.00027 | | | | | |
| | CMF / 0.01 - 2 mm | 0.01124 | 0.00053 | | | | | |
| | CMF / 2.01 - 4 mm | 0.00957 | 0.00043 | | | | | |
| | CMF / 4.01 - 6 mm | 0.00912 | 0.00038 | | | | | |
| | CMF / 6.01 - 8 mm | 0.00938 | 0.00039 | | | | | |
| | CMF / 8.01 - 10 mm | 0.01029 | 0.00051 | | | | | |
| | RMF / 0.01 - 2 mm | 0.01148 | 0.00075 | | | | | |
| | RMF / 2.01 - 4 mm | 0.01014 | 0.00025 | | | | | |
| | RMF / 4.01 - 6 mm | 0.00992 | 0.00024 | | | | | |
| | RMF / 6.01 - 8 mm | 0.01029 | 0.00026 | | | | | |
| | RMF / 8.01 - 10 mm | 0.01055 | 0.00038 | | | | | |
| Age x Brain Area x Distance | Younger / M1 / 0.01 - 2 mm | 0.01384 | 0.00121 | 16 | 528 | 2.278 | 0.052 | 0.016 |
| | Younger / M1 / 2.01 - 4 mm | 0.01293 | 0.00059 | | | | | |
| | Younger / M1 / 4.01 - 6 mm | 0.01224 | 0.00045 | | | | | |
| | Younger / M1 / 6.01 - 8 mm | 0.01267 | 0.00070 | | | | | |
| | Younger / M1 / 8.01 - 10 mm | 0.01316 | 0.00101 | | | | | |
| | Younger / S1 / 0.01 - 2 mm | 0.01291 | 0.00112 | | | | | |
| | Younger / S1 / 2.01 - 4 mm | 0.01070 | 0.00052 | | | | | |
| | Younger / S1 / 4.01 - 6 mm | 0.01068 | 0.00049 | | | | | |
| | Younger / S1 / 6.01 - 8 mm | 0.01235 | 0.00051 | | | | | |
| | Younger / S1 / 8.01 - 10 mm | 0.01370 | 0.00074 | | | | | |
| | Younger / SFG / 0.01 - 2 mm | 0.00844 | 0.00067 | | | | | |
| | Younger / SFG / 2.01 - 4 mm | 0.00784 | 0.00040 | | | | | |
| | Younger / SFG / 4.01 - 6 mm | 0.00766 | 0.00033 | | | | | |
| | Younger / SFG / 6.01 - 8 mm | 0.00736 | 0.00032 | | | | | |
| | Younger / SFG / 8.01 - 10 mm | 0.00727 | 0.00037 | | | | | |
| | Younger / CMF / 0.01 - 2 mm | 0.00919 | 0.00074 | | | | | |
| | Younger / CMF / 2.01 - 4 mm | 0.00776 | 0.00059 | | | | | |

| | | |
|---|---|---|
| Younger / CMF / 4.01 - 6 mm | 0.00757 | 0.00054 |
| Younger / CMF / 6.01 - 8 mm | 0.00785 | 0.00054 |
| Younger / CMF / 8.01 - 10 mm | 0.00845 | 0.00071 |
| Younger / RMF / 0.01 - 2 mm | 0.00849 | 0.00105 |
| Younger / RMF / 2.01 - 4 mm | 0.00895 | 0.00035 |
| Younger / RMF / 4.01 - 6 mm | 0.00903 | 0.00033 |
| Younger / RMF / 6.01 - 8 mm | 0.00977 | 0.00036 |
| Younger / RMF / 8.01 - 10 mm | 0.01051 | 0.00053 |
| Older / M1 / 0.01 - 2 mm | 0.02142 | 0.00125 |
| Older / M1 / 2.01 - 4 mm | 0.01834 | 0.00061 |
| Older / M1 / 4.01 - 6 mm | 0.01818 | 0.00046 |
| Older / M1 / 6.01 - 8 mm | 0.01987 | 0.00072 |
| Older / M1 / 8.01 - 10 mm | 0.02047 | 0.00104 |
| Older / S1 / 0.01 - 2 mm | 0.01657 | 0.00115 |
| Older / S1 / 2.01 - 4 mm | 0.01418 | 0.00054 |
| Older / S1 / 4.01 - 6 mm | 0.01368 | 0.00051 |
| Older / S1 / 6.01 - 8 mm | 0.01397 | 0.00053 |
| Older / S1 / 8.01 - 10 mm | 0.01483 | 0.00077 |
| Older / SFG / 0.01 - 2 mm | 0.01302 | 0.00069 |
| Older / SFG / 2.01 - 4 mm | 0.01110 | 0.00041 |
| Older / SFG / 4.01 - 6 mm | 0.01081 | 0.00034 |
| Older / SFG / 6.01 - 8 mm | 0.00993 | 0.00033 |
| Older / SFG / 8.01 - 10 mm | 0.00968 | 0.00038 |
| Older / CMF / 0.01 - 2 mm | 0.01329 | 0.00076 |
| Older / CMF / 2.01 - 4 mm | 0.01139 | 0.00061 |
| Older / CMF / 4.01 - 6 mm | 0.01068 | 0.00055 |
| Older / CMF / 6.01 - 8 mm | 0.01092 | 0.00055 |
| Older / CMF / 8.01 - 10 mm | 0.01212 | 0.00073 |
| Older / RMF / 0.01 - 2 mm | 0.01448 | 0.00108 |
| Older / RMF / 2.01 - 4 mm | 0.01132 | 0.00036 |
| Older / RMF / 4.01 - 6 mm | 0.01081 | 0.00034 |
| Older / RMF / 6.01 - 8 mm | 0.01082 | 0.00037 |
| Older / RMF / 8.01 - 10 mm | 0.01059 | 0.00055 |

**Supplementary Table 13. ANOVA results for pQSM values including the primary motor cortex (M1), primary somatosensory cortex (S1), superior frontal gyrus (SFG), caudal middle frontal cortex (CMF), and rostral middle frontal cortex (RMF).** Results for the ANOVA on pQSM values with factors age (n=18 younger adults, n=17 older adults), cortical depth (SF, OM, IM, DP), brain area (S1, M1, SFG, CMF, RMF) and venous distance (5 distances). Given are mean pQSM values in parts per million (Mean in ppm) and standard error of the mean (SEM), degrees of freedom of the numerator (DFn), degrees of freedom of the denominator (DFd), p-values (p), and effect sizes ($n_G^2$). P-values ≤ 0.05 are considered significant.

| | Conditions | Mean (ppm) | SEM (ppm) | Statistics | | | | |
|---|---|---|---|---|---|---|---|---|
| | | | | DFn | DFd | F | p | $\eta_G^2$ |
| Age | Younger | -0.00928 | 0.00032 | 1 | 33 | 42.594 | $2.066*10^{-7}$ | 0.192 |
| | Older | -0.01232 | 0.00033 | | | | | |
| Brain Area | M1 | -0.01051 | 0.00031 | 4 | 132 | 8.848 | $1.054*10^{-5}$ | 0.026 |
| | S1 | -0.01126 | 0.00027 | | | | | |
| | SFG | -0.01070 | 0.00024 | | | | | |
| | CMF | -0.01006 | 0.00037 | | | | | |
| | RMF | -0.01147 | 0.00024 | | | | | |
| Cortical depth | Deep | -0.01004 | 0.00027 | 3 | 99 | 78.915 | $1.510*10^{-13}$ | 0.154 |
| | Inner Middle | -0.00954 | 0.00029 | | | | | |
| | Outer Middle | -0.01061 | 0.00027 | | | | | |
| | Superficial | -0.01300 | 0.00028 | | | | | |
| Distance | 0.01 - 2 mm | -0.01283 | 0.00041 | 4 | 132 | 60.867 | $5.079*10^{-14}$ | 0.103 |
| | 2.01 - 4 mm | -0.01075 | 0.00023 | | | | | |
| | 4.01 - 6 mm | -0.01034 | 0.00019 | | | | | |
| | 6.01 - 8 mm | -0.01023 | 0.00022 | | | | | |
| | 8.01 - 10 mm | -0.00985 | 0.00025 | | | | | |
| Age x Distance | Younger / 0.01 - 2 mm | -0.01084 | 0.00056 | 4 | 132 | 3.339 | 0.049 | 0.006 |
| | Younger / 2.01 - 4 mm | -0.00940 | 0.00032 | | | | | |
| | Younger / 4.01 - 6 mm | -0.00903 | 0.00026 | | | | | |
| | Younger / 6.01 - 8 mm | -0.00879 | 0.00030 | | | | | |
| | Younger / 8.01 - 10 mm | -0.00834 | 0.00036 | | | | | |
| | Older / 0.01 - 2 mm | -0.01483 | 0.00058 | | | | | |
| | Older / 2.01 - 4 mm | -0.01210 | 0.00032 | | | | | |
| | Older / 4.01 - 6 mm | -0.01166 | 0.00027 | | | | | |
| | Older / 6.01 - 8 mm | -0.01167 | 0.00031 | | | | | |
| | Older / 8.01 - 10 mm | -0.01135 | 0.00037 | | | | | |
| Age x Brain Area | Younger / M1 | -0.00914 | 0.00043 | 4 | 132 | 1.922 | 0.122 | 0.006 |
| | Older / M1 | -0.00991 | 0.00038 | | | | | |
| | Younger / S1 | -0.00943 | 0.00033 | | | | | |
| | Older / S1 | -0.00825 | 0.00051 | | | | | |
| | Younger / SFG | -0.00967 | 0.00033 | | | | | |
| | Older / SFG | -0.01188 | 0.00045 | | | | | |
| | Younger / CMF | -0.01261 | 0.00039 | | | | | |
| | Older / CMF | -0.01196 | 0.00034 | | | | | |
| | Younger / RMF | -0.01187 | 0.00052 | | | | | |
| | Older / RMF | -0.01328 | 0.00034 | | | | | |
| Brain Area x Distance | M1 / 0.01 - 2 mm | -0.01009 | 0.00065 | 16 | 528 | 10.137 | $8.102*10^{-8}$ | 0.054 |
| | M1 / 2.01 - 4 mm | -0.01024 | 0.00028 | | | | | |
| | M1 / 4.01 - 6 mm | -0.01050 | 0.00023 | | | | | |
| | M1 / 6.01 - 8 mm | -0.01080 | 0.00030 | | | | | |
| | M1 / 8.01 - 10 mm | -0.01090 | 0.00049 | | | | | |
| | S1 / 0.01 - 2 mm | -0.01286 | 0.00056 | | | | | |
| | S1 / 2.01 - 4 mm | -0.01126 | 0.00032 | | | | | |
| | S1 / 4.01 - 6 mm | -0.01108 | 0.00024 | | | | | |
| | S1 / 6.01 - 8 mm | -0.01083 | 0.00027 | | | | | |
| | S1 / 8.01 - 10 mm | -0.01026 | 0.00037 | | | | | |
| | SFG / 0.01 - 2 mm | -0.01357 | 0.00050 | | | | | |
| | SFG / 2.01 - 4 mm | -0.01064 | 0.00027 | | | | | |
| | SFG / 4.01 - 6 mm | -0.01022 | 0.00021 | | | | | |
| | SFG / 6.01 - 8 mm | -0.00991 | 0.00020 | | | | | |
| | SFG / 8.01 - 10 mm | -0.00915 | 0.00025 | | | | | |
| | CMF / 0.01 - 2 mm | -0.01299 | 0.00082 | | | | | |
| | CMF / 2.01 - 4 mm | -0.01040 | 0.00037 | | | | | |
| | CMF / 4.01 - 6 mm | -0.00954 | 0.00034 | | | | | |
| | CMF / 6.01 - 8 mm | -0.00900 | 0.00035 | | | | | |
| | CMF / 8.01 - 10 mm | -0.00838 | 0.00036 | | | | | |
| | RMF / 0.01 - 2 mm | -0.01465 | 0.00062 | | | | | |
| | RMF / 2.01 - 4 mm | -0.01121 | 0.00025 | | | | | |
| | RMF / 4.01 - 6 mm | -0.01038 | 0.00023 | | | | | |
| | RMF / 6.01 - 8 mm | -0.01059 | 0.00026 | | | | | |
| | RMF / 8.01 - 10 mm | -0.01053 | 0.00035 | | | | | |
| Age x Brain Area x Distance | Younger / M1 / 0.01 - 2 mm | -0.00809 | 0.00091 | 16 | 528 | 0.614 | 0.672 | 0.003 |
| | Younger / M1 / 2.01 - 4 mm | -0.00876 | 0.00039 | | | | | |
| | Younger / M1 / 4.01 - 6 mm | -0.00931 | 0.00032 | | | | | |
| | Younger / M1 / 6.01 - 8 mm | -0.00958 | 0.00042 | | | | | |
| | Younger / M1 / 8.01 - 10 mm | -0.00996 | 0.00068 | | | | | |
| | Younger / S1 / 0.01 - 2 mm | -0.01125 | 0.00077 | | | | | |
| | Younger / S1 / 2.01 - 4 mm | -0.01016 | 0.00045 | | | | | |
| | Younger / S1 / 4.01 - 6 mm | -0.01005 | 0.00033 | | | | | |
| | Younger / S1 / 6.01 - 8 mm | -0.00939 | 0.00038 | | | | | |
| | Younger / S1 / 8.01 - 10 mm | -0.00871 | 0.00051 | | | | | |
| | Younger / SFG / 0.01 - 2 mm | -0.01193 | 0.00070 | | | | | |
| | Younger / SFG / 2.01 - 4 mm | -0.00953 | 0.00038 | | | | | |
| | Younger / SFG / 4.01 - 6 mm | -0.00924 | 0.00029 | | | | | |
| | Younger / SFG / 6.01 - 8 mm | -0.00878 | 0.00028 | | | | | |
| | Younger / SFG / 8.01 - 10 mm | -0.00768 | 0.00036 | | | | | |
| | Younger / CMF / 0.01 - 2 mm | -0.01088 | 0.00115 | | | | | |
| | Younger / CMF / 2.01 - 4 mm | -0.00902 | 0.00052 | | | | | |
| | Younger / CMF / 4.01 - 6 mm | -0.00779 | 0.00048 | | | | | |
| | Younger / CMF / 6.01 - 8 mm | -0.00718 | 0.00049 | | | | | |
| | Younger / CMF / 8.01 - 10 mm | -0.00640 | 0.00051 | | | | | |

| | | |
|---|---|---|
| Younger / RMF / 0.01 - 2 mm | -0.01203 | 0.00086 |
| Younger / RMF / 2.01 - 4 mm | -0.00954 | 0.00035 |
| Younger / RMF / 4.01 - 6 mm | -0.00878 | 0.00033 |
| Younger / RMF / 6.01 - 8 mm | -0.00903 | 0.00036 |
| Younger / RMF / 8.01 - 10 mm | -0.00895 | 0.00049 |
| Older / M1 / 0.01 - 2 mm | -0.01210 | 0.00093 |
| Older / M1 / 2.01 - 4 mm | -0.01172 | 0.00040 |
| Older / M1 / 4.01 - 6 mm | -0.01168 | 0.00033 |
| Older / M1 / 6.01 - 8 mm | -0.01203 | 0.00043 |
| Older / M1 / 8.01 - 10 mm | -0.01185 | 0.00070 |
| Older / S1 / 0.01 - 2 mm | -0.01447 | 0.00080 |
| Older / S1 / 2.01 - 4 mm | -0.01236 | 0.00046 |
| Older / S1 / 4.01 - 6 mm | -0.01212 | 0.00034 |
| Older / S1 / 6.01 - 8 mm | -0.01227 | 0.00039 |
| Older / S1 / 8.01 - 10 mm | -0.01182 | 0.00052 |
| Older / SFG / 0.01 - 2 mm | -0.01520 | 0.00072 |
| Older / SFG / 2.01 - 4 mm | -0.01174 | 0.00039 |
| Older / SFG / 4.01 - 6 mm | -0.01120 | 0.00030 |
| Older / SFG / 6.01 - 8 mm | -0.01104 | 0.00029 |
| Older / SFG / 8.01 - 10 mm | -0.01063 | 0.00037 |
| Older / CMF / 0.01 - 2 mm | -0.01510 | 0.00118 |
| Older / CMF / 2.01 - 4 mm | -0.01179 | 0.00053 |
| Older / CMF / 4.01 - 6 mm | -0.01129 | 0.00049 |
| Older / CMF / 6.01 - 8 mm | -0.01083 | 0.00050 |
| Older / CMF / 8.01 - 10 mm | -0.01035 | 0.00052 |
| Older / RMF / 0.01 - 2 mm | -0.01726 | 0.00089 |
| Older / RMF / 2.01 - 4 mm | -0.01289 | 0.00036 |
| Older / RMF / 4.01 - 6 mm | -0.01198 | 0.00033 |
| Older / RMF / 6.01 - 8 mm | -0.01216 | 0.00037 |
| Older / RMF / 8.01 - 10 mm | -0.01211 | 0.00050 |

**Supplementary Table 14. ANOVA Results for nQSM values including the primary motor cortex (M1), primary somatosensory cortex (S1), superior frontal gyrus (SFG), caudal middle frontal cortex (CMF) and rostral middle frontal cortex (RMF).** Results for the ANOVA on nQSM values with factors age (n=18 younger adults, n=17 older adults), cortical depth (SF, OM, IM, DP), brain area (S1, M1, SFG, CMF, RMF) and venous distance (5 distances). Given are mean nQSM values in parts per million (Mean in ppm) and standard error of the mean (SEM), degrees of freedom of the numerator (DFn), degrees of freedom of the denominator (DFd), p-values (p), and effect sizes ($n_G^2$). P-values $\leq$ 0.05 are considered as significant.
